# Cerebellar organoids model cell type-specific *FOXP2* expression during human cerebellar development

**DOI:** 10.1101/2024.12.20.628854

**Authors:** Elizabeth J. Apsley, Joey Riepsaame, Sally A. Cowley, Esther B. E. Becker

## Abstract

In this study, we demonstrate the potential of cerebellar organoids for studying features of early human cerebellar development. Forkhead box protein P2 (FOXP2) is a transcription factor associated with speech and language development that is highly expressed in the developing brain. However, little attention has been directed to the study of FOXP2 in the early developing cerebellum. We used CRISPR gene editing in human iPSCs to generate a fluorescent FOXP2-reporter line. By combining transcriptomic analysis of iPSC-derived cerebellar organoids with published cerebellar datasets, we describe the expression and identify potential downstream targets of FOXP2 in the early developing human cerebellum. Our results highlight expression of *FOXP2* in early human Purkinje cells and cerebellar nuclei neurons, and the vulnerability of these cell populations to neurodevelopmental disorders. Our study demonstrates the power of cerebellar organoids to model early human developmental processes and disorders.

## Introduction

The differentiation of human induced pluripotent stem cells (iPSCs) into cerebellar neurons and organoids has been shown to recapitulate early stages of cerebellar development^1–4^ and thus provides a human-specific model to probe the specification and maturation of cerebellar neurons. Single-cell RNA sequencing (scRNAseq) studies have profiled the cell types present in differentiated cerebellar organoids, including populations with characteristics of granule cells and Purkinje cells, the major populations of neurons in the cerebellum^4–6^. Moreover, following prolonged culture, iPSC-derived cerebellar organoids and plated neurons have been shown to develop spontaneous neuronal activity^3,4,6^. Following the establishment of robust methods, to date, iPSC-derived cerebellar models have been used to study the mechanisms of cerebellar diseases such as ataxia, medulloblastoma and pontocerebellar hypoplasia^7–10^. However, cerebellar organoid models have not yet been applied to investigating aspects of normal human cerebellar development.

In this study, we used iPSC-derived cerebellar organoids to investigate the development and properties of cells expressing the speech and language-associated gene *FOXP2* in the early human cerebellum. This gene encodes Forkhead box protein P2 (FOXP2), a transcription factor of the evolutionarily conserved FOXP subfamily. Over two decades ago, a mutation in *FOXP2* was identified as the monogenic cause of a severe speech and language disorder in a large multigenerational family^11^. However, despite considerable work, there is still relatively little understanding of the molecular mechanisms that underlie the function of FOXP2 and how this contributes to language development^12^. During development *FOXP2* is expressed across several brain regions including the cerebellum^13–15^. Interestingly, cerebellar alterations have been identified in neuroimaging studies of individuals affected by *FOXP2*-disorders and in genetic models of *Foxp2* disruption^13,16–18^. Homozygous disruption of *Foxp2* led to decreased size and foliation of the cerebellum, whilst heterozygous mice showed little or no gross cerebellar changes^19–22^. Cerebellar-specific knock-down of *Foxp2* in mice during embryonic development demonstrated the contribution of FOXP2 action in the cerebellum to normal brain function by reproducing some of the phenotype of global mutations, including impaired Purkinje cell dendritic development, gross motor deficits and altered vocalisation^23^. In contrast, postnatal Purkinje cell knock-out of *Foxp2* caused impaired performance only in more skilled motor tasks^24^. These differences in timing of *Foxp2* disruption and the severity of the resulting phenotype suggest that *Foxp2* has a particularly important role in newly differentiated Purkinje cells. However, studies of *Foxp2* disruption have largely focused on alterations in Purkinje cell dendritic development, without examining earlier stages of Purkinje cell differentiation or characterising other cerebellar cell types in detail.

Recent studies have highlighted important differences between mouse and human cerebellar development, including in the Purkinje cell lineage^25–27^. Human cerebellar development is highly protracted, and progenitor zones in the human developing cerebellum that give rise to the glutamatergic and GABAergic neuronal populations are more complex^25,27^. In addition, the developing human cerebellum exhibits significantly higher Purkinje cell abundances and a shift in the relative proportions of embryonic Purkinje cells subtypes compared to mouse^26^. These studies highlight a need for human-centric models to gain insights into mechanisms underlying early human cerebellar development.

To better understand the role of *FOXP2* in the specification and maturation of Purkinje cells in humans, we employed a human-specific cerebellar organoid model. We first showed that FOXP family genes *FOXP1, FOXP2* and *FOXP4* were robustly induced during organoid differentiation. We then generated a *FOXP2* reporter line and performed detailed characterisation of *FOXP2*-expressing cells in human cerebellar organoids. Transcriptomic analysis of these cells identified features of Purkinje cells and cerebellar nuclei neurons. *FOXP2*-positive neurons expressed a high number of genes associated with specific diseases including autism spectrum disorder (ASD). Our findings suggest that *FOXP2*-positive neurons in the developing human cerebellum are specifically vulnerable in neurodevelopmental disorders and demonstrate the potential of cerebellar organoids for investigating underlying disease mechanisms in distinct cell types.

## Results

### *FOXP* genes show distinct expression patterns in developing Purkinje cells

We first examined the expression of *FOXP2* in the developing human cerebellum using published transcriptomic datasets. Plotting *FOXP2* mRNA levels across human brain development, using the BrainSpan bulk RNA-sequencing dataset (https://www.brainspan.org/), confirmed particularly high *FOXP2* expression in the cerebellum, striatum and dorsal thalamus in the first and second trimester (Fig. 1A). By birth, *FOXP2* expression in the cerebellar cortex decreased substantially (Fig. 1A). This time period coincides with a shift in cell type proportions in the cerebellum due to a massive expansion of granule cells and therefore relative decrease in the Purkinje cell population^26,28^. Within the cerebellum, single-nucleus RNA-sequencing (snRNA-seq) data from the prenatal human cerebellum confirmed strong *FOXP2* expression in Purkinje cells from 8 post conception weeks (PCW) (Fig. 1B)^26^. *FOXP2* expression remained high in Purkinje cells up to 20 PCW, but was decreased at birth consistent with the bulk RNA-sequencing data (Fig. 1A, B)^26^. The low number of Purkinje cells captured in postnatal samples limited characterisation of expression at later timepoints.

**Figure 1.**
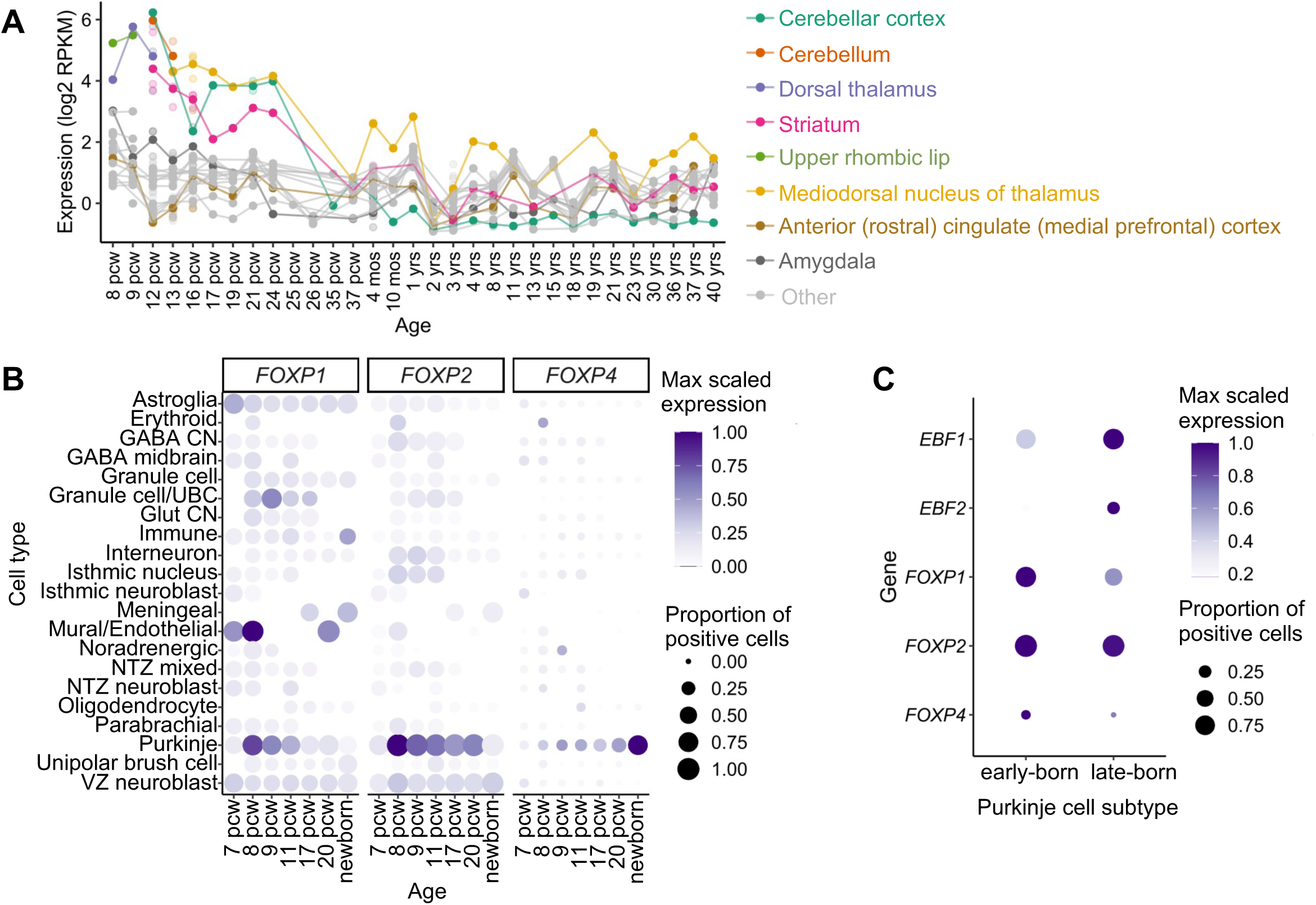
*FOXP2* is expressed highly in developing human Purkinje cells. **(A)** *FOXP2* is highly expressed in the cerebellum during human brain development. Bulk RNA-seq data from BrainSpan (https://www.brainspan.org/). Regions with highest expression are depicted in colour, with other brain regions shown in grey. Where multiple samples were present for a timepoint, the mean value is a solid circle. Individual samples are represented by open circles. pcw, post-conception weeks; mos, months after birth; yrs, years after birth. **(B)** Expression of *FOXP* genes in the developing human cerebellum plotted across timepoints and cell types based on published snRNA-seq data ^26^. Point size indicates the proportion of expressing cells in each cluster and the colour scale shows average expression levels. Samples with <10 cells were excluded. **(C)** *FOXP* genes show differential expression across Purkinje cell subtypes with *FOXP1* marking early-born Purkinje cells (FDR=4.58 x 10^-302^, log2foldchange, log2FC=0.613). *FOXP2* (FDR=1.82 x 10^-22^, log2FC=0.158) and *FOXP4* (FDR=8.51 x 10^-20^, log2FC=0.0436) are slightly enriched in the early-born Purkinje cell population, in contrast to late-born markers *EBF1* (FDR=2.16 x 10^-304^, log2FC=0.725) and *EBF2* (FDR=5.40 x 10^-165^, log2FC=0.302). Analysis based on snRNAseq data from Sepp et al., 2024^26^.

Dimerization of FOXP2 is essential for DNA-binding and transcriptional regulation^29^. FOXP2 can form both homodimers or heterodimers with the closely related FOXP1 or FOXP4, with combinations differentially regulating target genes^29,30^. Therefore, the combinations of other FOXP proteins co-expressed with FOXP2 are significant to its function. To consider the possible FOXP protein interactions in the developing human cerebellum, we also analysed the expression of *FOXP1* and *FOXP4* in transcriptomic data of the developing human cerebellum. Both *FOXP1* and *FOXP4* were enriched in the Purkinje cell cluster and thus overlapping with *FOXP2*, though with different trajectories (Fig. 1B). Similar to *FOXP2*, *FOXP1* expression peaked at 8 PCW and decreased over time, however *FOXP1* showed an earlier fall, with Purkinje cell expression dropping by 17 PCW (Fig. 1B). In contrast, *FOXP4* expression increased at later stages of prenatal development, with highest expression occurring in the newborn cerebellum (Fig. 1B). Similar trends in *Foxp* gene expression were found in the developing mouse cerebellum (Fig. S1A). *Foxp*2 and *Foxp4* were most highly expressed in Purkinje cells, with levels of *Foxp2* peaking in development and levels of *Foxp4* increasing later in adult Purkinje cells as previously described (Fig. S1A)^15,31,32^. Expression of *Foxp1* in mouse Purkinje cells was relatively low overall, but as in human peaked early in embryonic development (Fig. S1A).

Purkinje cells are known to be a heterogenous population and can be grouped into subtypes on the basis of gene expression and location from early embryonic development^33–35^. In mouse, the differential expression of *Foxp1* and *Foxp2* has been suggested to contribute to defining embryonic Purkinje cell subpopulations, with multiple studies defining a subtype exhibiting high *Foxp1* expression^26,36^. To investigate whether this also applies in human development, we compared *FOXP* gene expression across early- and late-born human Purkinje cell subtypes^26^. *FOXP2* showed expression across both subtypes although we found a small but significant enrichment in the early-born subtype (False discovery rate, FDR = 1.8 x10^-22^, log2FC = 0.158). *FOXP1* and *FOXP4* expression was strongly enriched in the early-born Purkinje cell subtype, and little expression was found in the late-born Purkinje cells (*FOXP1* FDR = 4.58 x10^-302^, log2FC = 0.613, *FOXP4* FDR = 8.51 x10^-20^, log2FC = 0.0436) (Fig. 1C). We also observed similar expression of *Foxp* genes across mouse Purkinje cell subtypes. *Foxp2* was expressed evenly across embryonic Purkinje cell subtypes, with a small significant enrichment in the *Foxp1*-high subtype (Enrichment in each cluster: *Cdh9*-subtype FDR = 1, *Etv1*-subtype FDR = 1, *Foxp1*-subtype FDR = 0.0185, *Rorb*-subtype FDR = 1) (Fig. S1B). *Foxp1* and *Foxp4* were differentially expressed across mouse Purkinje cell subtypes, with both highly expressed in the early-born *Foxp1-*labelled cluster (Enrichment in *Foxp1*-cluster, *Foxp1* FDR = 1.2 x10^-83^, *Foxp4* FDR = 4.07 x10^-6^) (Fig. S1B). Together, these findings suggest that FOXP heterodimers are likely to be present in developing human Purkinje cells with varying composition across subtypes and over time.

Having observed similar patterns in *FOXP* expression in the snRNAseq data between mouse and human, we used mouse cerebellar sections to examine how the transcriptomic variation related to the spatial distribution of FOXP proteins. In agreement with the relatively even expression across subtypes in the transcriptomic data, FOXP2 protein expression was observed broadly in Purkinje cells. By E15.5 FOXP2 was detected in Purkinje cells migrating from the ventricular zone to form the Purkinje cell plate visible at E18.5, and later in the Purkinje cell layer (Fig. S1C-F). The only exception we observed was in the most posterior lobule in lateral sections, consistent with previous descriptions of little or no FOXP2 in a small population of Purkinje cells in the most lateral, rostral and ventral region of the cerebellum^14^ (Fig. S1E). Reflecting the differential expression that we had identified across transcriptional defined subtypes, FOXP1 and FOXP4 showed more heterogenous expression in Purkinje cells at E15.5 (Fig. S1C), with FOXP1 only present in more lateral sections. At E18.5, FOXP1-positive cells were largely found on the outer surface of the cerebellum, and no longer overlapped with FOXP2-expressing Purkinje cells within the cerebellum (Fig. S1D). FOXP4 continued to display differential expression between Purkinje cell clusters at E18.5 (Fig S1D), however, in the postnatal mouse cerebellum, FOXP4 expression was found uniformly across all Purkinje cells (Fig. S1F). Together, the gene expression and imaging data confirmed the expression of FOXP2 in developing human and mouse Purkinje cells, and indicate that FOXP heterodimers are likely to be present with varying composition across subtypes and over time. Our observations also suggest that FOXP4, in addition to previously described FOXP1, shows differential expression across embryonic Purkinje cell subtypes in both mouse and human.

### iPSC-derived cerebellar organoids provide a human model to study *FOXP* expression in the developing cerebellum

Our analysis of transcriptomic data from the developing human cerebellum confirmed enriched expression of FOXP2 in developing human Purkinje cells and suggested differential expression of *FOXP* genes may contribute to Purkinje cell subtype patterning. Whilst these patterns could be validated in mouse tissue, we next sought a human model to investigate the function of FOXP2 in the developing human cerebellum. Cerebellar organoids provide an attractive *in vitro* system to study early human cerebellar development, modelling the specification of cerebellar identity and production of cerebellar neurons from both the rhombic lip and ventricular zone lineages^1,5^. To investigate whether cerebellar organoids could be used to investigate FOXP2 in the developing human cerebellum, we first characterised the expression of *FOXP* family genes throughout differentiation in iPSC-derived cerebellar organoids. Cerebellar organoids were generated from human iPSCs according to our established protocol, using FGF2 and insulin to direct differentiation of stem cell aggregates to cerebellar identity^37^(Fig. 2A). By day 35 of the differentiation protocol, organoids contain cells from both cerebellar germinal zones shown by immunostaining against KIRREL2 labelling the ventricular zone, and PAX6 labelling rhombic lip-derived granule cell progenitors and granule cells, indicating successful differentiation^37^ (Fig. S2). mRNA expression of all three *FOXP* genes was found to be strongly upregulated within the first 21 days of cerebellar differentiation (Fig. 2B). *FOXP2* and *FOXP4* mRNA expression plateaued from day 49 of differentiation, whilst *FOXP1* mRNA expression continued to increase (Fig. 2B). Similar trends in protein expression were confirmed by immunostaining. FOXP2 protein was occasionally detected in organoids at day 35, but expression was consistently seen in organoids at day 49 or later (Fig. 2C). FOXP1 and FOXP4 protein expression was detectable from day 21 (Fig. 2C). To predict the potential FOXP dimers present in cerebellar organoids, we quantified the co-expression of FOXP proteins at day 63, a timepoint with high FOXP2 expression. FOXP2 was commonly co-expressed with other FOXP proteins: of cells staining positive for FOXP2, 75% were also positive for FOXP1 and 63% were FOXP4-positive (Fig. 2D-F). Together, our analysis of cerebellar organoid samples showed strong induction of *FOXP2* during cerebellar differentiation and indicates that expression of *FOXP* genes in the early developing cerebellum is recapitulated using iPSC-derived cerebellar organoids. The presence of both double and single FOXP-positive cells suggests the presence of different Purkinje cell subpopulations in the cerebellar organoids as described *in vivo*^26,36^.

**Figure 2.**
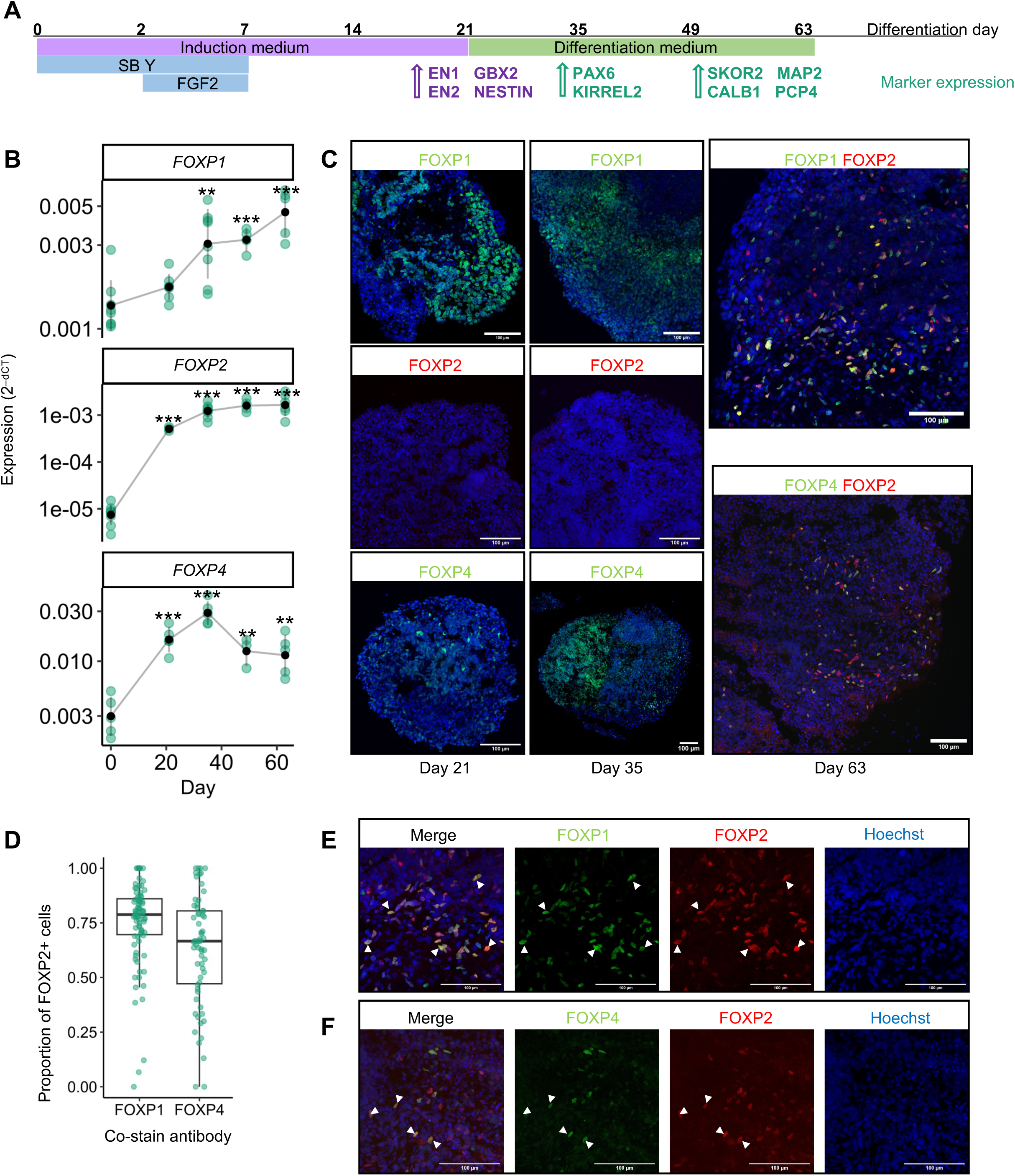
Cerebellar organoids recapitulate expression of *FOXP* family genes during differentiation. **(A)** Schematic of cerebellar organoid differentiation. Human iPSC aggregates are patterned towards cerebellar identity using growth factors including Fibroblast growth factor 2 (FGF2), TGFβ inhibitor SB431542 (SB), anti-apoptotic factor rock inhibitor, Y-27632 (Y) and insulin (component of Induction medium). Specific cell lineage markers used to monitor differentiation at corresponding timepoints are indicated. **(B)** *FOXP* genes significantly increase in expression during cerebellar organoid development. Expression calculated as 2^-dCT^, relative to housekeeper genes *ACTB* and *GAPDH*. Significant change in expression compared to day 0 determined using a t-test on dCT values. \**p*<0.05, \*\**p*<0.01, \*\*\**p*<0.001. Green points represent individual differentiations (n=3-9), black points show mean and error bars representing mean±SD. **(C)** Imaging of FOXP protein expression across cerebellar organoid differentiation. FOXP1 and FOXP4 (green) can be consistently detected by immunostaining from day 21, and FOXP2 (red) from day 49. Nuclei were stained with Hoechst (blue). Scale bars = 100 µm. **(D)** The colocalization of FOXP protein staining was quantified as the proportion of FOXP2-positive cells co-expressing FOXP1 and FOXP4 respectively. Data points represent separate images, with more than three sections imaged per organoid, and organoids from five differentiation experiments. **(E)** Higher magnification images showing co-expression of FOXP1 (green) and **(F)** FOXP4 (green) with FOXP2 (red), as marked by arrowheads. Nuclei were stained with Hoechst (blue). Scale bars = 100 µm.

### Generation of a *FOXP2* iPSC reporter line facilitates live visualisation and isolation of *FOXP2*-positive cells in cerebellar organoids

Cerebellar organoids do not recapitulate the spatial organisation of the developing cerebellum and therefore showed a heterogenous distribution of FOXP2-expressing cells across the organoid (Fig. 2C) compared to the distinctive staining pattern observed in tissue sections the developing mouse cerebellum (Fig. S1C-F). This lack of spatial organization makes it hard to identify specific cerebellar cell types purely by their position in the organoid and typically requires analysis of marker gene expression in fixed samples. To facilitate the study of *FOXP2*-expressing cells in cerebellar organoids, including live visualising this population, we generated an iPSC line containing a fluorescent reporter for *FOXP2* expression. We targeted the endogenous *FOXP2* locus to achieve an accurate readout of *FOXP2* expression. mNeonGreen (hereafter mNeon) was selected for bright fluorescent properties given the relatively weak expression of an endogenous promoter^38^. CRISPR-mediated gene editing was employed using a single-stranded DNA donor, encoding a self-cleaving peptide P2A^39,40^ and the fluorescent protein mNeon flanked by homology arms (Fig. 3A). iPSC clones with successful homozygous integration of the mNeon sequence were screened for by PCR (Fig. 3B, C & Fig. S3A-C), and accurate insertion of the reporter gene was confirmed by Sanger sequencing (Fig. S3D). We selected two clones (A6B, B2D) with homozygous insertion of mNeon into the *FOXP2* locus for future experiments.

**Figure 3.**
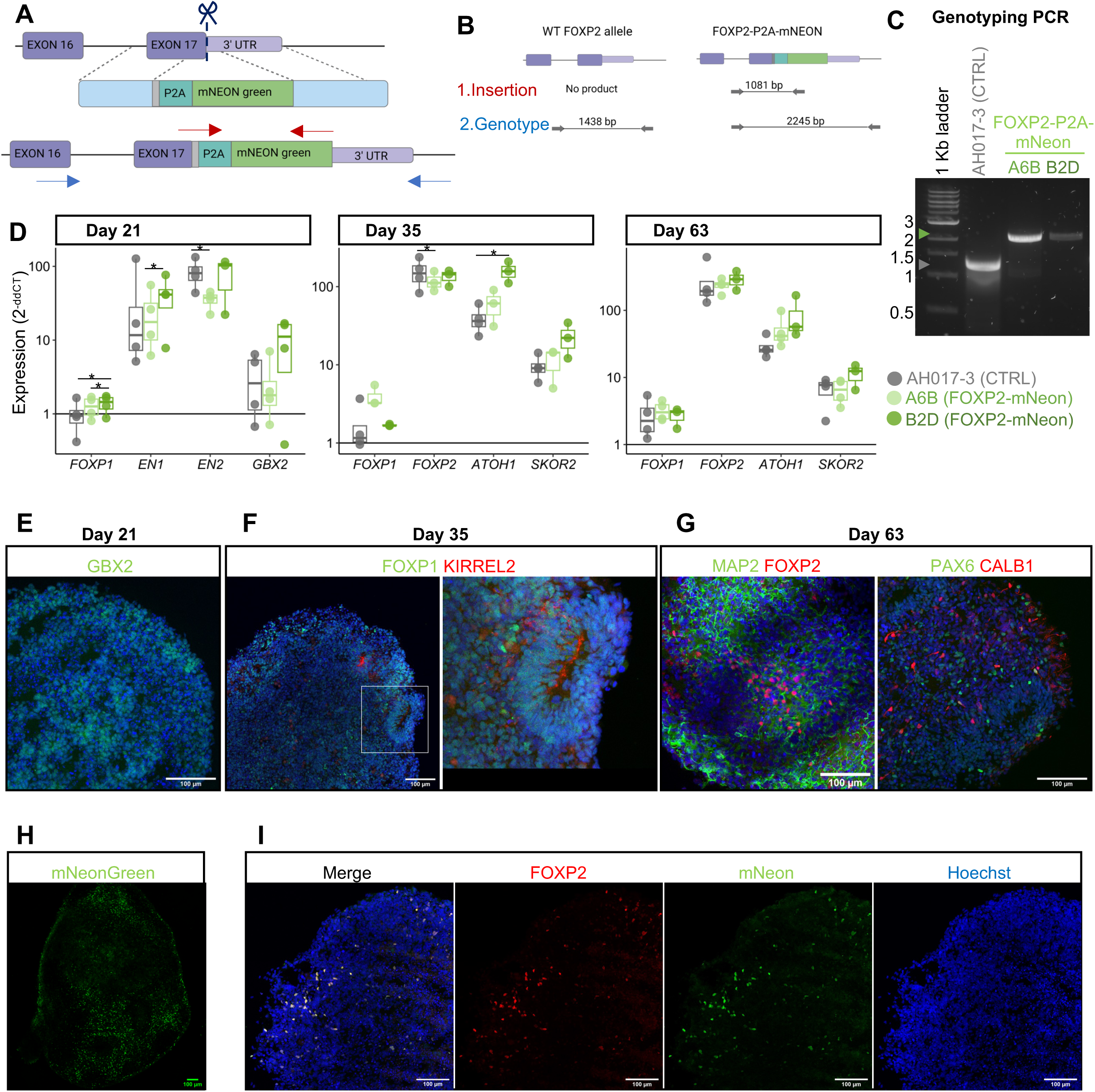
Generation of FOXP2-mNeon iPSCs and cerebellar organoids. **(A)** CRISPR strategy for the generation of a FOXP2-mNeon reporter line. A specific gRNA targets Cas9 (scissors) to the endogenous *FOXP2* stop codon (immediately upstream of the 3’ untranslated region (UTR)). The donor repair construct contains a P2A-mNeon sequence surrounded by homology arms (light blue) matching the target sequences indicated by the dashed lines. The result of homology-dependent repair shown below is a FOXP2-P2A-mNeon reporter gene. The position of primers used for screening is indicated by arrows. **(B)** Schematic of PCR products resulting from screening clones, first detecting insertion and then genotyping. **(C)** Products of the genotyping PCR on a 1% agarose gel from the parental AH017-3 line and two clones with mNeon insertion (A6B & B2D) demonstrate successful gene editing. **(D)** Expression of cerebellar markers at days 21, 35 and 63 of cerebellar organoid differentiation using two FOXP2-mNeon clones (A6B, light green & B2D, dark green) and the parental line (AH017-3, grey) by RT-qPCR. Expression calculated relative to iPSCs (day 0) as 2^-ddCT^. Dots indicate individual differentiations (n = 3-4). Significant differences in gene expression between lines at each timepoint were tested for using an ANOVA followed by post-hoc Tukey’s HSD. \**p*<0.05, \*\**p*<0.01, \*\*\**p*<0.001. **(E)** Expression of hindbrain marker GBX2 (green) at day 21 of cerebellar differentiation using FOXP2-mNeon clone B2D. Nuclei were stained with Hoechst (blue). Scale bar = 100 µm **(F)** Expression of markers of the cerebellar ventricular zone (FOXP1, green & KIRREL2, red) at day 35 of cerebellar differentiation using FOXP2-mNeon clone B2D. Nuclei were stained with Hoechst (blue). Scale bars = 100 µm. **(G)** Expression of markers of Purkinje cells (FOXP2, red & CALB1, red), Granule neurons (PAX6, green) and mature neurons (MAP2, green) at day 63 of cerebellar differentiation using FOXP2-mNeon clone B2D. Nuclei were stained with Hoechst (blue). Scale bars 100 µm**. (H)** Live-cell imaging of mNeon fluorescence in a day 58 cerebellar organoid generated from FOXP2-mNeon clone B2D, showing distribution of FOXP2-positive cells. **(I)** Immunofluorescent staining of FOXP2 (red) and mNeon (green) confirms accurate co-expression in sections of a day 63 cerebellar organoid generated using FOXP2-mNeon clone B2D. Nuclei were stained with Hoechst (blue). Scale bars = 100 µm.

No mutations resulting from the CRISPR process were detected upon sequencing of the top six high-risk off-target sites predicted by online tools CCTop (https://cctop.cos.uniheidelberg.de:8043/) and CRISPOR (http://crispor.tefor.net/) for the gRNA used (Fig. S4). In addition, FOXP2-mNeon clones showed equivalent gene dosage to control iPSC AH017-3 by SNP array (noting a loss of heterozygosity without copy number loss on Chr17p in the parental AH017-3 working stock (Fig. S5)). FOXP2-mNeon clones displayed typical iPSC morphology and showed widespread expression of pluripotency markers NANOG and TRA-1-60 when assessed by flow cytometry (Fig. S6A, B). Moreover, the generated reporter line iPSCs could be differentiated successfully into all three germ lineages following respective directed differentiation, confirming their pluripotency (Fig. S6C). Together these results indicate that the generated iPSC reporter line maintained normal stem cell characteristics following the CRISPR editing and clone selection process.

Next, we tested the capability of the newly generated *FOXP2* reporter line to differentiate reliably into cerebellar organoids. Both FOXP2-mNeon clones successfully generated cerebellar organoids with typical growth, morphology and induction of cerebellar genes used for quality control (Fig. 3D). Markers of the midbrain-hindbrain boundary were present by day 21, with significant upregulation of *EN1* and *EN2* (t-test on dCT values, day 21 vs iPSC, p-values: A6B - *EN1 =* 1.30 x 10^-4^*, EN2* = 6.16 x 10^-6^*, GBX2* = 0.114, *FOXP1* = 0.102. B2D – *EN1 =* 6.43 x 10^-4^*, EN2 =3.23* x 10^-4^*, GBX2* = 0.0303, *FOXP1* = 0.0825) (Fig. 3D) and positive staining for GBX2 (Fig. 3E). At day 35 and 63, reporter line-derived cerebellar organoids expressed markers for progenitors and neurons of both key cerebellar neuron lineages: granule cells (*ATOH1, PAX6*) and Purkinje cells (*KIRREL2, SKOR2, CALB1*) (Fig. 3D, F, G). For both FOXP2-mNeon clones, there was significant induction of Purkinje cell (*FOXP1, FOXP2, SKOR2*) and granule cell (*ATOH1*) marker genes, at day 35 and day 63 compared to in iPSCs as measured by qPCR (t-test on dCT values, day 35 vs iPSC or day 63 vs iPSC, p<0.01 for all genes) (Fig. 3D). Organoids generated from both FOXP2-mNeon reporter line clones showed very similar expression of profiled markers compared to the parental line (Fig. 3D). At day 63, organoids widely expressed MAP2 indicating a high proportion of differentiated neurons (Fig. 3G). Following generation of FOXP2-mNeon cerebellar organoids, mNeon expression could be successfully visualised by live-cell fluorescent microscopy (Fig. 3H). Immunofluorescent staining of fixed organoid sections showed tight co-expression of the mNeon reporter with FOXP2 (Fig. 3I), verifying the accuracy of mNeon signal as a readout of endogenous FOXP2 expression. The subcellular localisation of mNeon was observed as nuclear in both live-cell microscopy and immunostaining (Fig. 3H, I). This suggested that the fluorescent mNeon tag was not separated from the nuclear-localized FOXP2 by the intended P2A-mediated cleavage. The presence of a FOXP2-mNeon fusion protein, instead of separated FOXP2 and mNeon proteins, was confirmed by western blotting (Fig. S7). Together, these results demonstrate that the generated FOXP2-mNeon iPSC lines could be differentiated reliably to cerebellar identity, including a *FOXP2*-positive population accurately labelled live by mNeon fluorescence.

### Transcriptomic analysis of *FOXP2*-expressing cells in cerebellar organoids reveals Purkinje cell and cerebellar nuclei neuron identities

To further characterise the cellular identity of *FOXP2*-positive (*FOXP2*+) cells in the cerebellar organoids, we performed RNA-sequencing on sorted populations of mNeon-positive and mNeon-negative cells from differentiated *FOXP2* reporter line organoids. FOXP2-mNeon cerebellar organoids were dissociated to single cells at day 63 of differentiation, around the peak of FOXP2-positive cells detected by immunostaining (Fig. 2C). Flow cytometry was used to sort live cells into mNeon-positive and -negative populations, with ∼3% of live cells in the mNeon-positive gate (Fig. S8).

Principal component analysis revealed clear separation of the mNeon-positive and -negative samples around principal component 1 (PC1), with this parameter responsible for 95% variance compared to a very small effect between individual differentiations (Fig. 4A). Differential expression analysis confirmed the isolation of the *FOXP2*-expressing cells in the mNeon-positive population (Fig. 4B). In total, 7644 genes were identified as differentially expressed below the adjusted *p*-value threshold p<0.05, with 3655 genes increased and 3989 decreased in the mNeon-positive population relative to the mNeon-negative cells. In agreement with our immunostaining results (Fig. 2C, D), *FOXP1* and *FOXP4* expression was significantly enriched in the mNeon-positive, *FOXP2*-expressing population (Fig. 4B).

**Figure 4.**
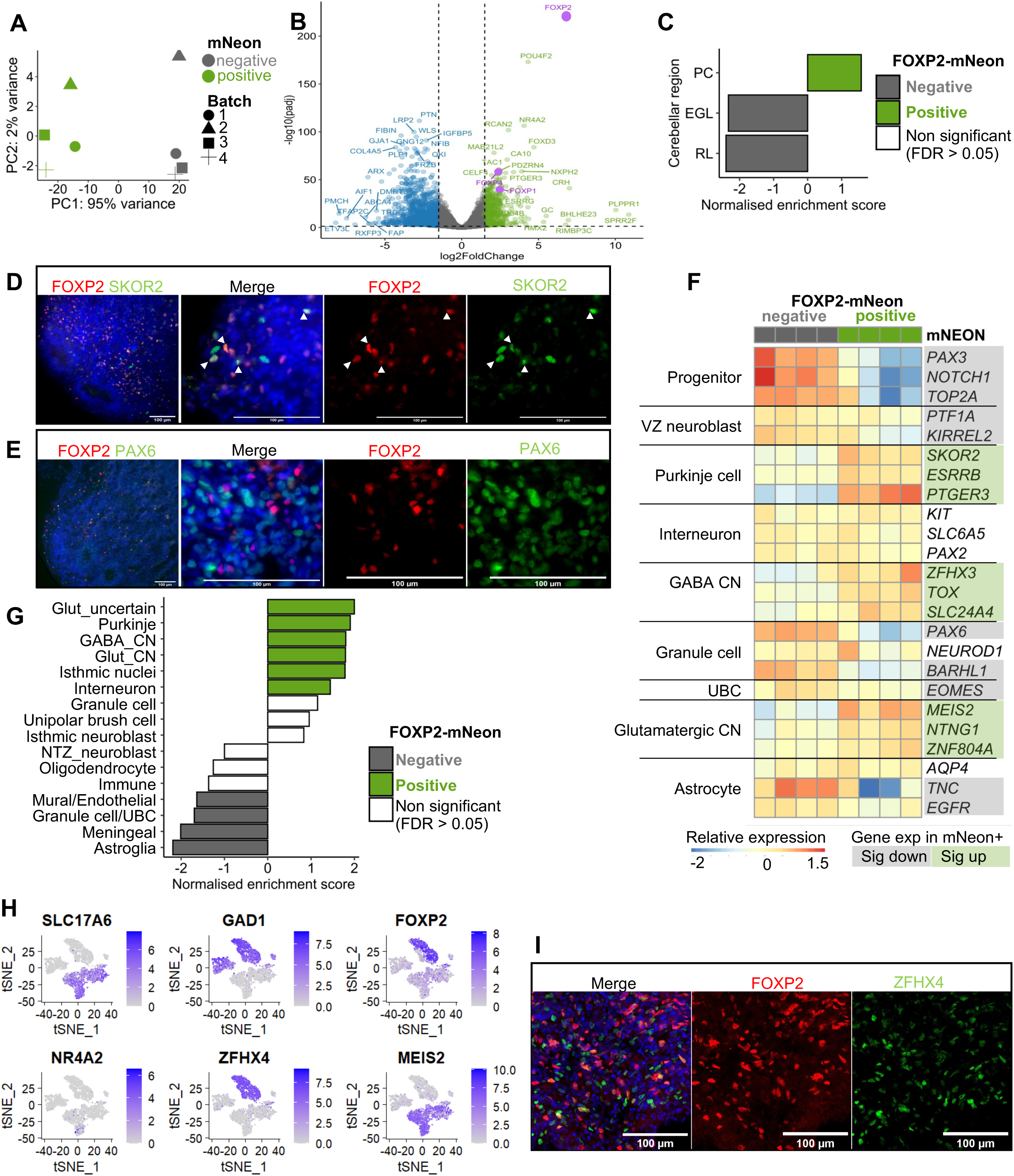
Transcriptomic characterisation of mNeon-sorted populations identifies *FOXP2* expression in Purkinje cells and cerebellar nuclei neurons in cerebellar organoids. **(A)** mNeon-positive (green) and mNeon-negative (grey) populations showed clear separation across the x axis, with principal component 1 (PC1) accounting for the majority of variance between samples. Individual differentiations shown by different shapes contribute a small amount of variance across PC2 (y axis). **(B)** Differentially expressed genes in the mNeon-positive populations are displayed on volcano plot. Genes increased in mNeon-positive samples are shown in green (log2FoldChange (log2FC)>1.5, *p*adj<0.05) include *FOXP2* (log2FC=6.82, *p*adj=1.75×10^-224^), *FOXP1* (log2FC=2.4, *p*adj=5.2×10^-41^) and *FOXP4* (log2FC=2.3, *p*adj=1.61 x10^-58^). Genes significantly decreased in mNeon-positive samples are shown in blue (log2FC<-1.5, *p*adj<0.05). Differential gene expression Wald test with Benjamini-Hochberg multiple testing adjustment using DESeq2. **(C)** Enrichment of genes differentially expressed between fetal human cerebellar regions (Aldinger et al., 2021^28^) in FOXP2-mNeon-postive and -negative cerebellar organoid populations. Gene set enrichment analysis (GSEA) calculated a normalised enrichment score and FDR for each geneset. PC: Purkinje cell layer; EGL: external granule layer; RL: rhombic lip. **(D)** Immunostaining for FOXP2 (red) with Purkinje cell marker SKOR2 (green) using day 63 cerebellar organoid sections. Arrowheads indicate co-expression. Nuclei were stained with Hoechst (blue). Scale bar = 100 µm. **(E)** Immunostaining for FOXP2 (red) with granule cell marker PAX6 (green) of cerebellar organoid sections (day 49) did not identify co-expression. Nuclei were stained with Hoechst (blue). Scale bar = 100 µm. **(F)** Expression of specific cell type markers across organoid populations indicated that the *FOXP2*+ cerebellar organoid cells contain a mixture of Purkinje cell and cerebellar nuclei (CN) neurons. Colour scale shows expression scaled for each gene. Genes highlighted have *p*adj<0.05 and are increased (green, log2FC>0) or decreased (grey, log2FC<0) in the *FOXP2*+ population. VZ: ventricular zone; UBC: unipolar brush cells. **(G)** The distribution of genes specifically expressed in distinct cerebellar cell types was assessed across cerebellar organoid using GSEA. Genesets significantly enriched (FDR<0.05) in the *FOXP2*+ or *FOXP2-* cerebellar organoid population are coloured in green and grey, respectively. CN: cerebellar nuclei; Glut: glutamatergic; NTZ: nuclear transitory zone; UBC unipolar brush cell **(H)** Plotting published snRNAseq data from human cerebellar nuclei neurons shows their division into glutamatergic excitatory (*SLC17A6*) and GABAergic inhibitory (*GAD1*) clusters (Kebschull et al., 2020^41^). *FOXP2* expression is enriched in an *GAD1+* inhibitory cluster, but also shows broader expression. Expression of selected genes upregulated in the *FOXP2*+ cerebellar organoid population (*NR4A2, ZFHX4, MEIS2*) are shown across the same dataset. **(I)** Immunostaining of day 63 cerebellar organoid, using antibodies against *FOXP2* (red) and *ZFHX4* (green). Nuclei were stained with Hoechst (blue). Scale bar = 100 µm.

To assess the cell-type identity most closely represented by the *FOXP2*+ population, we compared the distribution of marker genes across the flow cytometry-sorted cerebellar organoid samples. Gene set enrichment analysis (GSEA) was performed using differentially expressed genes in fetal human cerebellar layers^28^. The *FOXP2*+ population showed a significant enrichment for genes distinguishing the Purkinje cell layer (PCL), whilst the *FOXP2*-population showed significant enrichment for genes highest in the rhombic lip (RL) and external granule cell layer (EGL) (Fig. 4C). These results are consistent with *FOXP2* as a marker of Purkinje cells and suggest that the remaining organoid population contained proliferative progenitors, including granule cell progenitors, and non-neuronal cells. Immunostaining confirmed overlap in protein expression of the Purkinje cell-specific marker SKOR2 within the FOXP2+ population and exclusion of expression of PAX6, a marker of granule cell precursors and neurons (Fig. 4D, E). Following the identification of Purkinje cell features in the FOXP2+ cerebellar organoid population, we next tested whether there was any Purkinje cell subtype enrichment within the *FOXP2*+ organoid population using markers to distinguish between the early-born and late-born clusters^26^. Comparison of differentially expressed genes across the organoid populations identified a slight enrichment of late-born Purkinje markers in the FOXP+ population (Fisher’s exact test, *p*=0.016, data not shown).

To refine the identity of the *FOXP2*+ cerebellar organoid population further, we examined expression of selected markers of different cerebellar cell types (Fig. 4F). The cell type specificity of these marker genes was first demonstrated by plotting expression across cell types in recent snRNA-seq data profiling of the developing cerebellum^26^ (Fig. S8). The *FOXP2*+ population showed significantly increased expression of markers of Purkinje cells, particularly those present in earlier stages of cerebellar development (*e.g*., *SKOR2, ESRRB, PTGER3, LHX1, LHX5, TFAP2A*). Genes associated with progenitors (*PAX3, NOTCH1, TOP2A*) and ventricular zone neuroblast (*KIRREL2*, *PTF1A*) were significantly decreased in the *FOXP2*+ population, indicating maturity of the *FOXP2*+ population as differentiated neurons in contrast to progenitors. Other cerebellar cell lineages were excluded from the *FOXP2*+ population, based on decreased expression of markers for granule cells (*PAX6, ATOH1, BARHL1*) and unipolar brush cells (*EOMES, LMX1A*), and no significant difference in markers for GABAergic interneurons (*PAX2, KIT, SLC6A5*). However, several genes associated with both glutamatergic (*SLC17A6, MEIS2, NTNG1, ZNF804A*), and GABAergic (*ZFHX3, ZFHX4, TOX, SLC24A4*) neurons found in the cerebellar nuclei (CN) were also found to be enriched in the *FOXP2*+ organoid population. To expand from our short curated list of cell type markers, we calculated differentially expressed genes for each cluster present in the Sepp et al., 2024^26^ dataset to determine a list of genes specific to each cerebellar cell type, and these marker gene lists were used in GSEA. In support of our initial conclusions, the *FOXP2*+ population showed significant enrichment for markers of Purkinje cells (Fig. 4G). GSEA also identified an upregulation of markers of GABAergic and glutamatergic CN, and to a lesser extent interneurons, in the *FOXP2*+ population (Fig. 4G). In addition, there was significant enrichment for markers of isthmic nuclei neurons (genes including *NR4A2* and *SCG2*) and a cluster labelled as “glutamatergic_uncertain” (genes including *VSNL1, RIMS1, FSTL5*), both clusters from the nuclear transitory zone lineage, which gives rise to glutamatergic CN and isthmic nuclei neurons^26^.

Our results suggest that CN neurons expressing *FOXP2* were amongst the *FOXP2*+ population in cerebellar organoids. Supporting the expression of *FOXP2* in CN, we also detected sparse FOXP2-expressing cells in the cerebellar white matter in mouse immunostained cerebellar sections (Fig. S1F), in line with previous descriptions^15,31^. Therefore, we sought to determine whether *FOXP2* is also expressed in the human CN, and in which cell types. We made use of a published snRNAseq dataset of human CN neurons^41^. The CN contain excitatory (primarily glutamatergic) and inhibitory (primarily GABAergic) neurons that can be distinguished by expression of *SLC17A6* and *GAD1* respectively^41^ (Fig. 4H). We found that expression of *FOXP2* was highest in a cluster of inhibitory CN neurons, annotated as “inferior olive projection neurons”, but was also widely expressed at a lower level across excitatory CN clusters (Fig. 4H). Next, we plotted the expression of genes enriched in the *FOXP2*+ organoid population associated with the isthmic nuclei (*NR4A2*), GABAergic CN neurons (*ZFHX4*) and glutamatergic CN neurons (*MEIS2*) in the human CN dataset. *ZFHX4* showed a similar profile to *FOXP2* with enrichment in the same inhibitory neuron cluster, whilst *MEIS2* and *NR4A2* were higher in excitatory neurons (Fig. 4H). Immunostaining was performed to confirm the co-expression of one of these CN markers, ZFHX4 with FOXP2 in cerebellar organoids (Fig. 4I). Together, these results confirmed that the presence of markers of CN neurons in the *FOXP2*+ cells of cerebellar organoids is reflective of the multiple cell types which express FOXP2 in the human cerebellum, most notably Purkinje cells and CN.

### *FOXP2*-positive cells show enrichment for pathways relating to neuronal activity

FOXP2 acts as a transcriptional regulator, capable of transcriptional repression or activation, depending on interacting partners^29,42,43^ and therefore influencing a range of cellular processes. To investigate potential downstream pathways of FOXP2 in the developing human cerebellum, we performed GSEA using biological process gene ontology (GO) terms. GSEA identified an enrichment for terms related to synaptic processes, neuronal activity and signal transduction in the *FOXP2*+ population (Fig. 5A). These pathways included upregulation of genes such as *GLRA*2, *CELF4*, *GRIN3A* and *GABRA3* for regulation of membrane potential, and synaptic vesicle components including synaptotagmins (*SYT7, SYT12, SYT5*), synaptophysin (*SYP*), synapsin (*SYN1*) and synaptic vesicle glycoproteins (*SV2C, SV2A*). Terms identified for the *FOXP2*-population were more varied and related to extracellular matrix, cytoplasmic translation and connective tissue development (Fig. 5B). Our approach can only identify pathways upregulated in *FOXP2*+ cells and cannot distinguish which are directly regulated by FOXP2. However, previous studies in non-cerebellar cells have identified similar pathways related to neurogenesis and synaptic signalling regulated by FOXP2^44–47^. Therefore, our data suggest a probable contribution of FOXP2 to these processes in cerebellar development.

**Figure 5.**
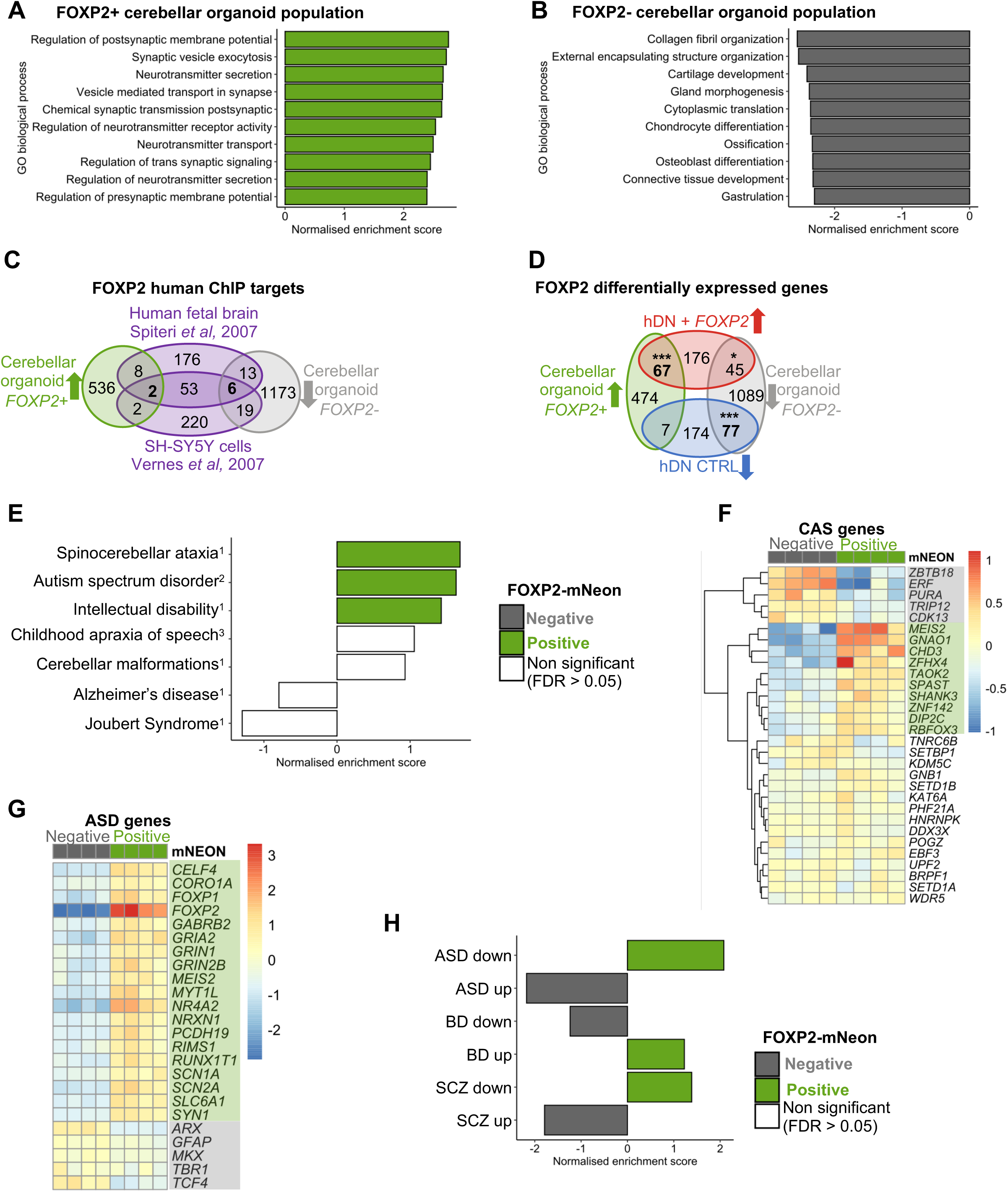
The *FOXP2*+ population in cerebellar organoids is enriched for expression of pathways relating to synaptic activity and for genes associated with certain neurodevelopmental disorders. **(A)** Upregulated and **(B)** downregulated biological processes in the *FOXP2*+ cerebellar organoid population as calculated by GSEA. Normalised enrichment scores for the top 10 gene ontology (GO) biological process terms are shown. **(C)** Differentially expressed genes between *FOXP2*+ (green) and *FOXP2*-(grey) cerebellar organoid populations were compared to FOXP2 targets identified in published FOXP2-ChIP studies on neuronal samples (Vernes et al., 2007^45^ & Spiteri et al., 2007^44^) and overlaps shown. **(D)** Differentially expressed genes were compared between *FOXP2*+ (green, left) and *FOXP2*-(grey, right) cerebellar organoid populations and human differentiated neurons with *FOXP2* (red, top) and without *FOXP2* (blue, bottom) expression (Hickey et al., 2019^43^). Significance of geneset overlap was determined using a Fisher’s exact test. \**p*<10^-5^, \*\*\**p*<10^-26^ **(E)** GSEA identified an enrichment in genes associated with spinocerebellar ataxia, ASD and intellectual disability in the *FOXP2*+ cerebellar organoid population. Genesets were collated from multiple sources: ^1^previous testing in human cerebellar scRNAseq (Aldinger et al., 2021^28^). ^2^SFARI class 1 risk genes (https://gene.sfari.org/). ^3^published CAS genes (Kaspi et al. 2022^54^; Eising et al. 2019^55^; Hildebrand et al. 2020^56^). Bars are coloured according to significance (FDR<0.05) and direction of enrichment. **(F)** Expression of CAS-associated genes across *FOXP2*-positive and -negative cerebellar organoid populations. Genes significantly differentially expressed (*p*adj<0.05) between FOXP2-sorted populations are highlighted in green (increased) or grey (decreased). **(G)** Expression of ASD-associated genes which showed significant differential expression (*p*adj<0.05) between *FOXP2*-positive and -negative cerebellar organoid populations. Genes highlighted according to change in the *FOXP2*-positive population in green (increased) or grey (decreased). **(H)** Enrichment of genes differentially expressed in post-mortem brain samples from individuals with autism spectrum disorder (ASD), bipolar disorder (BD) and schizophrenia (SCZ). Differentially expressed genes identified by Gandal et al., 2018^70^, genes which were significantly increased in affected individuals compared to healthy controls were classified as ‘up’, and those decreased as ‘down’. GSEA was performed to calculate a normalised enrichment score and FDR for each geneset across *FOXP2*-positive and -negative organoid populations.

In order to focus on transcriptional changes, which could be directly mediated by FOXP2 as opposed to differences in cell type identity, genes differentially expressed across organoid populations were compared to known FOXP2 targets. Several published studies have identified FOXP2 targets using chromatin immunoprecipitation (ChIP). However, there is very limited overlap in the results of these studies, and thus currently no consensus list of FOXP2-regulated genes^12^. As no FOXP2-ChIP studies have been performed on isolated cerebellar tissue, we focused on comparison with human neuronal studies, using human fetal brain^44^ and a human neuroblastoma cell line SH-SY5Y stably expressing *FOXP2*^45^. When selecting cerebellar organoid differentially expressed genes for this analysis, we applied a stricter threshold of an absolute log2FoldChange in expression greater than 1.5 and adjusted *p*-value below 0.05 to focus on the genes with most specific expression to one population. This yielded a list containing 578 genes increased and 1253 genes decreased in the *FOXP2*+ population. From the two FOXP2-ChIP studies, only 61 common targets were identified, two of these were significantly increased (*FOXD3, PDZRN*) in the *FOXP2*+ population in cerebellar organoids and six were found to be significantly decreased (*PTN, LRP2, COL4A5, HTRA1, LRP4, NOTCH2*) (Fig. 5C). We also examined the expression of a broader list of genes shown to be downstream of FOXP2-regulation in human neurons in the cerebellar organoids. These genes were identified by differential gene expression analysis of neurons differentiated *in vitro* from human neural progenitors which do not express *FOXP2* (CTRL) or with exogenous *FOXP2* expression^43^. Interestingly, a significant number of the genes present in both datasets showed the same pattern in expression when comparing between *FOXP2*+ and *FOXP2*-cells: 67 genes were significantly upregulated in the *FOXP2*+ vs *FOXP2*-cells of cerebellar organoids and in *FOXP2*+ human differentiated neurons vs control human differentiated neurons, and 77 genes were decreased in both (Fig. 5D). These results suggest that FOXP2 regulates a common set of genes in neuronal differentiation.

### Genes associated with neurodevelopmental disorders are enriched in *FOXP2*+ cells

Having explored pathways active in normal development, we next sought to identify whether the *FOXP2*+ cerebellar population was likely to be affected in disease by examining the expression of disease-associated genes across FOXP2-positive and -negative cerebellar organoid populations. Aberrant cerebellar development is linked to a range of neurodevelopmental disorders (NDD) including autism spectrum disorder (ASD)^48–50^. Moreover, developmental abnormalities also likely contribute to cerebellar diseases that are classically regarded as neurodegenerative conditions such as spinocerebellar ataxia (SCA)^51–53^. Employing GSEA using gene sets for cerebellar disorders, as previously examined in human fetal cerebellum^28^, and a list of CAS-risk genes^54–56^, we found a significant enrichment for genes associated with specific disorders including SCA, ASD and intellectual disability (ID) in the *FOXP2*+ cerebellar organoid population (Fig. 5E). There was also an enrichment, albeit non-significant, for CAS-associated genes as well as cerebellar malformations (Fig. 5E). As expected, no enrichment was identified in genes associated with Alzheimer’s disease, a neurodegenerative disease not linked to cerebellar development, and Joubert Syndrome, a neurodevelopmental ciliopathy. Joubert Syndrome genes have been previously shown to be enriched in the ciliated cell population of the cerebellar organoids^5^.

Childhood apraxia of speech (CAS) is the core phenotype of the *FOXP2*-related speech and language disorder^57^. More recently, additional monogenic causes of CAS have been identified in a series of sequencing studies of affected probands and their families, offering more insight into the gene networks regulating speech^54–56^. We further examined the expression of individual CAS genes in *FOXP2*+ cerebellar cells to identify co-expressed genes which might act in cooperation with *FOXP2* in the early cerebellum. Five genes were significantly decreased, and ten genes were significantly increased in the *FOXP2*+ population (*p*adj<0.05) (Fig. 5F). Interestingly, CAS genes with increased expression included several genes expressed in the CN such as *MEIS2, ZFHX4* and *RBFOX3* (Fig. 5F).

Whilst *FOXP1* has been repeatedly associated with ASD, including the presence of ASD-symptoms within the definition of FOXP1-syndrome^58^, the association of *FOXP2* and ASD has been less certain, with both positive and negative reports^13^. Interestingly, we found that ASD-associated genes which were differentially expressed between the *FOXP2*+ and *FOXP2*-organoid populations included *FOXP1* as well as several genes involved in neuronal activity, particularly synaptic transmission (*GABRB1, GRIA2, GRIN1, GRIN2B, SCN1A, SCN2A, SLC6A1*) (Fig. 5G). Given the strong enrichment for ASD associated genes in the *FOXP2*+ cerebellar organoid population, we also examined cerebellar organoid expression of genes with altered expression in ASD, using an additional dataset profiling differentially expressed genes in post-mortem brains of individuals with neurological disorders^70^ (Fig. 5H). We found that genes that were decreased in individuals with ASD were significantly enriched in the *FOXP2*+ organoid population, whereas genes that were upregulated in ASD post-mortem brains were enriched in the *FOXP2*-population in the cerebellar organoids (Fig. 5H). Interestingly, genes increased in individuals with bipolar disorder and gene decreased in individuals with schizophrenia were also significantly enriched in the *FOXP2*+ organoid population, whilst genes decreased in bipolar and genes increased in schizophrenia were enriched in the *FOXP2*-organoid population (Fig. 5H).

Together, our findings suggest that *FOXP2*+ cell populations might be particularly vulnerable to distinct NDDs and highlight specific gene networks associated with this vulnerability. Moreover, our results underscore the utility of the cerebellar organoid model to further study the disease mechanisms in vulnerable cerebellar cell types these disorders.

## Discussion

In this work, we used cerebellar organoids to study the expression of *FOXP2* in the developing human cerebellum. We successfully generated a fluorescent iPSC reporter line to visualise *FOXP2* expression live. Differentiation of this FOXP2-mNeon iPSC line into cerebellar organoids facilitated detailed characterisation of the *FOXP2*+ population from amongst the many organoid cell types. We identified features of Purkinje cells and CN neurons in the *FOXP2*+ cerebellar organoid population by comparison to published snRNAseq human cerebellar datasets. The *FOXP2*+ cerebellar population showed enrichment for molecular pathways related to neurogenesis and synaptic signalling. Moreover, we identified enrichment of a significant number of disease-associated genes in the *FOXP2*+ cerebellar organoid cells. Together, our results profile *FOXP2*+ cells in the developing human cerebellum and highlighted these cell types as important for understanding multiple NDDs. We also demonstrate the potential of cerebellar organoids to model early human cerebellar development and disease.

Cerebellar organoids provided a reliable, human-specific system to examine the expression of FOXP proteins across early cerebellar development. The expression of *FOXP1*, *FOXP2* and *FOXP4* in the human cerebellum and in cerebellar organoids showed similarities to previous descriptions of *Foxp* gene expression in mouse development^15,31,32^. Our analysis found *FOXP2* widely expressed across embryonic human Purkinje cells, with a slight enrichment in the early-born subtype. In comparison, scRNAseq from embryonic mouse cerebellum identified differences in level of *Foxp2* across Purkinje cell subtypes^36^. We saw greater variation in the distribution of *FOXP1*, with *FOXP1* expression significantly higher in the human early-born Purkinje cell subtype, mirroring multiple studies in mouse and human identifying high *Foxp1* in early-born lateral Purkinje cell clusters^26,36^. In comparison to other *FOXP* genes, *FOXP4* was expressed in a much lower proportion of fetal human Purkinje cells, but again showed higher expression in the human early-born Purkinje cell subtype. We also observed a distinct spatial distribution of FOXP4 across Purkinje cells in the developing mouse cerebellum and found an enrichment in the early-born, *Foxp1*-high subtype of mouse embryonic Purkinje cells. Our findings suggest that, similar to FOXP1, FOXP4 might contribute to defining embryonic Purkinje cell subtypes, a role which has not previously been reported. In cerebellar organoids, we found that FOXP2 was frequently co-expressed with other FOXP proteins, but also identified the presence of single FOXP-positive cells. We therefore hypothesise that cerebellar organoids may contain multiple Purkinje cell subtypes reflective of *in vivo* developmental processes, and so may allow further investigation into the developmental processes and physiological phenotypes of different human Purkinje cell subtypes.

Our findings draw attention to the expression of *FOXP2* in human CN neurons, and the recapitulation of this cell type-specific expression in cerebellar organoids. Histological studies in mouse have previously reported FOXP2 expression in the CN, however the focus of FOXP2 functional studies has largely been on Purkinje cells^14,15,31^. Along with *FOXP2*, we found that additional genes identified as monogenic causes of CAS such as *ZFHX4* are expressed in the same human CN populations. The CN act to transmit information out from the cerebellum to other brain regions, and also have an important feedback role in the inferior olive-cerebellar inhibitory circuits^53,59^. Our findings warrant further research into the role of these neurons in studies of FOXP2 disruption and NDDs more generally.

Beyond speech and language disorders, we found enrichment for genes associated with other NDDs in the *FOXP2*+ cerebellar organoid population. Our findings in the human cerebellar organoids are consistent with an enrichment of SCA and ASD genes observed in Purkinje cell clusters of human developing cerebellum^28^, underscoring the power of the cerebellar organoids in modelling relevant developing human cell populations that are vulnerable in disease. In particular, ASD-risk genes and genes downregulated in ASD post-mortem brains were enriched in the *FOXP2*+ population, whilst genes upregulated in ASD were lower in the *FOXP2*+ population. This result may reflect a decreased population of specific cell types, such as Purkinje cells or other GABAergic neurons sharing similar expression profiles or an altered phenotype such as reduced neuronal activity in ASD. Excitatory-inhibitory imbalance is a proposed mechanism underlying ASD and has been investigated in the context of the cerebral cortex and its organoid models, but not the cerebellum^49,60^. We propose that cerebellar organoids offer a promising model to understand the mechanisms of different genetic contributors to ASD during relevant developmental windows. Notably, the strong expression of the ASD-associated *FOXP1* gene in specific human Purkinje cell subtypes and CN neurons calls for further investigation into the function of FOXP1 in these cell types during normal development as well as NDDs.

In this study, we have shown that cerebellar organoids can recapitulate cell type populations present in the developing human cerebellum, including the presence of a *FOXP2*+ population expressing markers of Purkinje cells and CN. Whilst published snRNAseq datasets from human *in vivo* samples provided a useful tool to validate our model, these resources provide only static snapshots of development. In contrast, cerebellar organoids provide an *in vitro* human-model to dynamically test mechanisms of cell-type development and disease in the cerebellum.

## Resource availability

### Lead contact

Requests for further information and resources should be directed to lead contact Esther Becker.

### Materials availability

Cell lines used in this study are available from the lead contact, but may require a completed Materials Transfer Agreement and reasonable compensation by requestor for its processing and shipping.

### Data and code availability

RNA sequencing data has been deposited to GEO (GSE285003). Code is available at https://github.com/LizzyApsley/CerebellumFOXP2

## Acknowledgements

We would like to thank Vincent Cheng and Dianali Rodriguez Fernandez for assisting with cryosectioning, and Mohamed Ibrahim for extracting mouse cerebellar samples. We would also like to thank Robert Hedley and Vasiliki Tsioligka for providing technical assistance in sorting organoid populations at The Don Mason Facility of Flow Cytometry, Sir William Dunn School of Pathology, University of Oxford, and David Sims for aligning the bulk RNA sequencing data. Library preparation and sequencing was performed by the Oxford Genomics Centre, University of Oxford. The Dunn School Bioimaging Facility, University of Oxford, provided confocal imaging facilities.

This work was supported by funding from UKRI-BBSRC [BB/M011224/1].

## Author contributions

Conceptualization: E.J.A and E.B.E.B; Experimentation and analysis: E.J.A; CRISPR editing design: J.R; Stem cell culture and CRISPR-mediated gene editing technical expertise: S.A.C; Writing – original draft: E.J.A and E.B.E.B; Writing – review and editing: E.J.A., J.R., S.A.C. and E.B.E.B.; Funding acquisition: E.B.E.B.

## Methods

### Key resources table

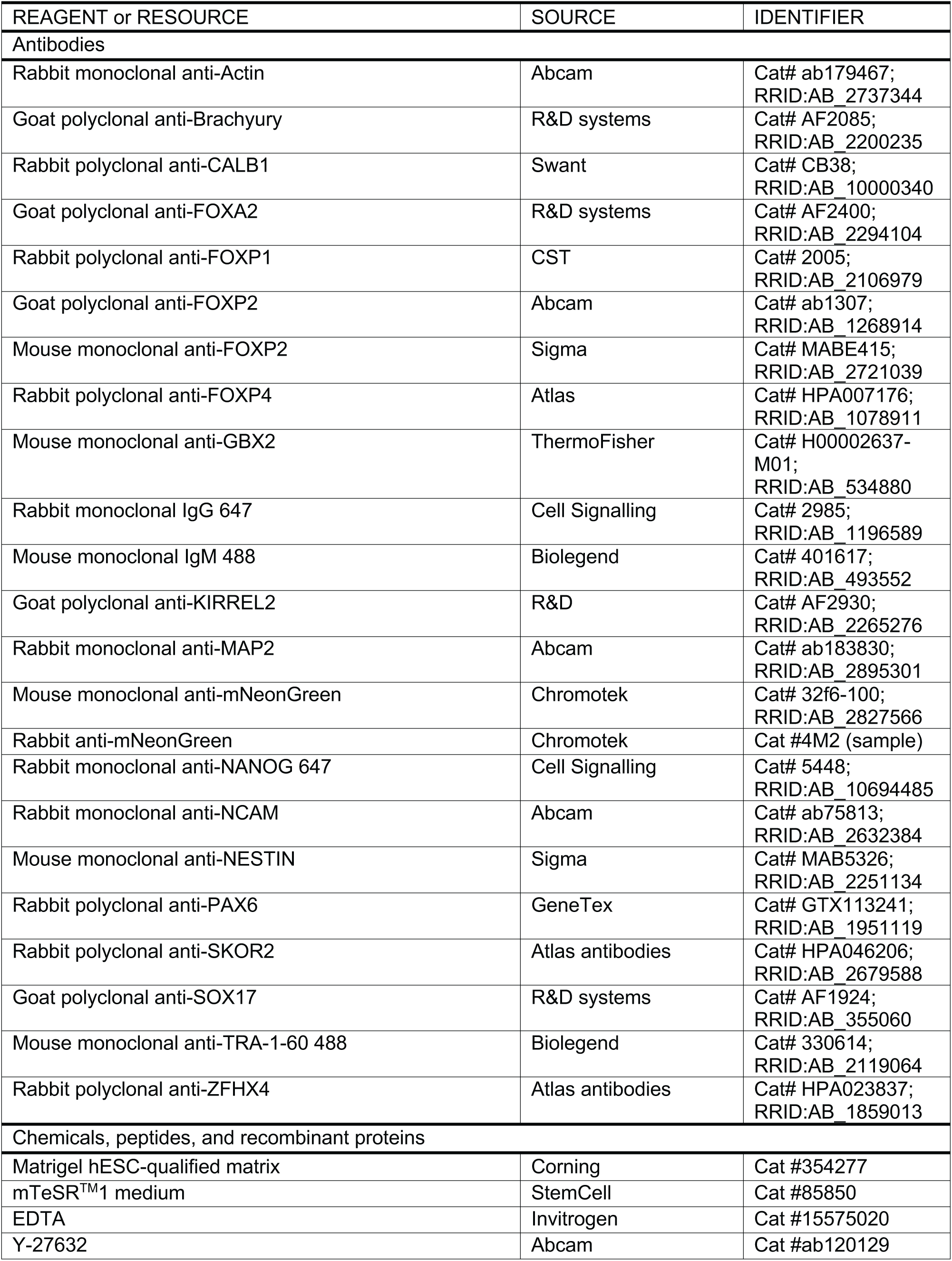

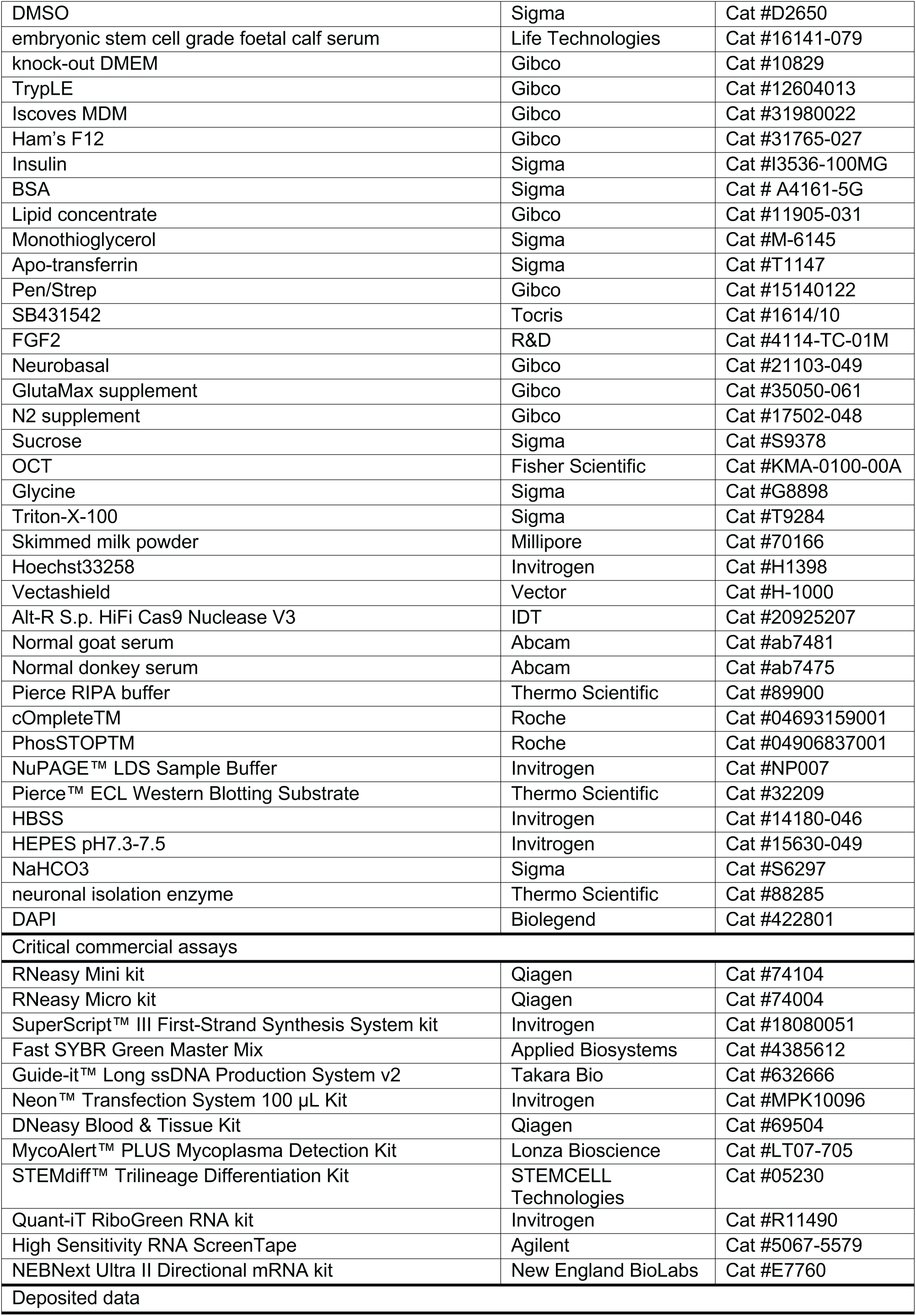

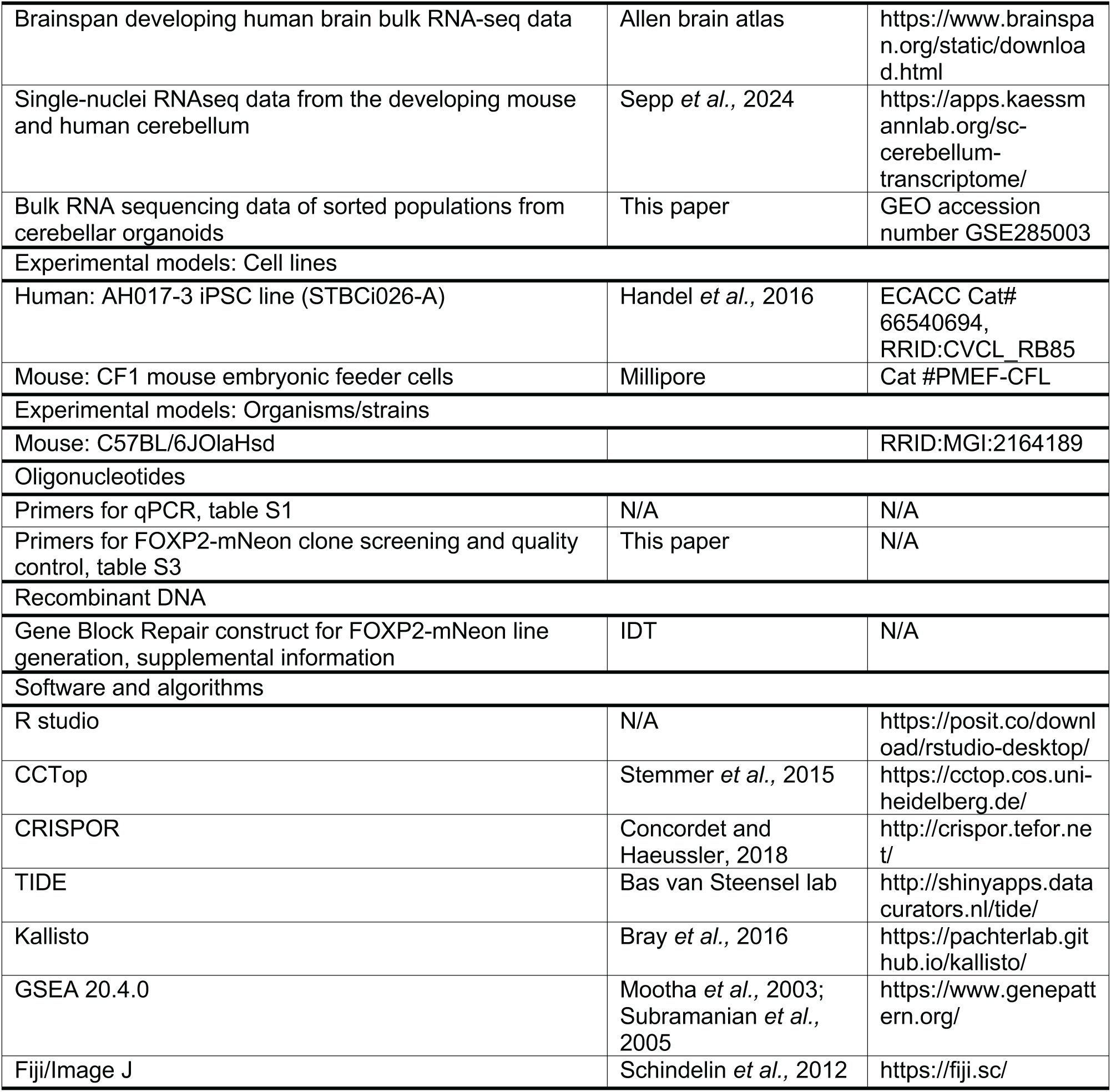

### Experimental model and study participant details

#### iPSC culture

Human iPSCs were grown on Matrigel hESC-qualified matrix (Corning, 354277) and fed daily with mTeSR^TM^1 medium (StemCell, 85850). Wells were passaged with 0.5mM EDTA (Invitrogen, 15575020) when 70-80% confluent. Rock inhibitor (10 μM Y-27632, Abcam, ab120129) was added to mTeSR^TM^1 medium to promote survival for the first 24 hour after thawing, subsequent culture and passaging was performed without Y-27632. iPSCs were expanded to generate a working stock and frozen in freezing medium (10% DMSO (Sigma, D2650), 30% embryonic stem cell grade foetal calf serum (ES-FCS, Life Technologies, 16141-079), 60% knock-out DMEM (Gibco, 10829)) at 2 x 10^6^ cells/vial. iPSCs were passaged at least once post thaw before differentiation. No antibiotics were used in iPSC culture and cells were visibly inspected for contamination to ensure sterility. We used the control iPSC line AH017-3 (STBCi026-A, ECACC Cat# 66540694, RRID:CVCL_RB85), derived from skin fibroblasts from a female donor by sendai virus reprogramming^61^. This line has been previously characterised and used for differentiation to cerebellar organoids^5,61^.

### Methods details

#### Analysis of published datasets

Human developing brain bulk RNA-seq data was downloaded from Brainspan (https://www.brainspan.org/static/download.html Accessed 06.01.2023). Processed single-nuclei RNAseq data from developing human cerebella was downloaded from https://apps.kaessmannlab.org/sc-cerebellum-transcriptome/ ^26^. The data was loaded in R and packages SingleCellExperiment (SingleCellExperiment_1.20.0), tidyverse (tidyverse_2.0.0) and reshape2 (reshape2_1.4.4), ggplot2 (ggplot2_3.4.3) and RColorBrewer (RColorBrewer_1.1-3) were used for generating plots.

#### Differentiation of human cerebellar organoids

Cerebellar organoids were generated from iPSCs following an established protocol^37^. In brief, iPSCs were dissociated using 1x TrypLE (Gibco, 12604013) and reaggregated in 96-well ultra-low attachment V-bottom plates (Greiner Bio-One, 651970) at a concentration of 1.25 x 10^4^ cells per well. Aggregates were seeded in Induction medium (50% Iscoves MDM (Gibco, 31980022), 50% Ham’s F12 (Gibco, 31765-027), 7 μg/ml insulin (Sigma, I3536-100MG), 5 mg/ml BSA (Sigma, A4161-5G), 1x Lipid concentrates (Gibco, 11905-031), 450 μM Monothioglycerol (Sigma, M-6145), 15 μg/ml Apo-transferrin (Sigma, T1147), 1x Pen/strep (Gibco, 15140122)), supplemented with 50 μM Y-27632 (Abcam, ab120129) and 10 μM SB431542 (Tocris, 1614/10). On day two, 20 μl of medium was removed and replaced with 20 μl of Induction medium with Y-27632 and SB431542 and 250 ng/ml FGF2 (R&D, 4114-TC-01M), resulting in a concentration of 50 ng/ml FGF2 in each well. On day seven, a 1/3 volume medium change was performed using Induction medium with no additional supplements. On day 14, organoids were transferred to 48-well ultra-low attachment plates (Greiner Bio-One, 677970) in 200 μl of fresh Induction medium per well. After 21 days, medium was switched to Differentiation medium (Neurobasal (Gibco 21103-049), 1x GlutaMax (Gibco 35050-061), 1x N2 (Gibco 17502-048), 1x Pen/Strep (Gibco, 15140122)) and a complete medium change performed every 7 days up to day 35. Beyond day 35, organoids were kept in suspension with Differentiation medium volume increased to 300 μl/well and a 1/3-1/2 volume medium change performed three times per week.

#### RT-qPCR

RNeasy Mini or Micro kits (Qiagen, 74104 or 74004) were used for isolation of RNA from iPSC pellets and cerebellar organoids. Around 20-30 organoids were sufficient for RNA extraction at day 21, 15-20 at day 35 and 8-10 at later timepoints. For each reaction, 0.5-1 μg of RNA was used for generating cDNA with the SuperScript™ III First-Strand Synthesis System kit (Invitrogen, 18080051). cDNA was then diluted 1:10 for use in RT-qPCR. Primers (Table S1) were tested using a standard curve to determine efficiency and melt-curves checked for a single qPCR product peak. RT-qPCR was performed using Fast SYBR Green Master Mix (Applied Biosystems, 4385612) on StepOne plus Reverse transcription PCR system (Applied Biosciences). Each gene-sample pair was performed in triplicate, with the average value taken from these technical replicates. Expression of genes of interest was calculated using the dCT method relative to housekeeper genes *ACTIN* and *GAPDH*. No significant changes in the expression of housekeeper genes were found across the differentiation time course. Statistical tests were performed on dCT values.

#### Immunostaining

Organoids were fixed in 4% PFA for 15-20 min at room temperature, washed once in PBS and then transferred into 20% sucrose (Sigma, S9378) to equilibrate overnight at 4°C. Next, organoids were embedded in Optimal Cutting Temperature compound (OCT, Fisher Scientific, KMA-0100-00A) and frozen for cryo-sectioning.

Before staining, slides were warmed to room temperature and hydrated in PBS for 10 min. Sections were treated with 0.1 M glycine (Sigma, G8898) for 30 min, permeabilised with 0.3% Triton-X-100 (Sigma, T9284) in PBS (PBST) for 30 min, and blocked in 2% milk (Millipore, 70166) in PBST for 1 hour. Primary antibodies (Table S2) were incubated in block overnight at 4°C. After three washes with 0.05% Tween in PBS, secondary antibodies (1:1000, Alexa Fluor, Invitrogen) were incubated for 1 hour. Wash steps were repeated and nuclei stained with 1 μg/ml Hoechst33258 (Invitrogen, H1398) in PBS before mounting in Vectashield (Vector, H-1000). Slides were imaged using a Zeiss Axioplan2 upright fluorescent microscope or a Zeiss-880 laser scanning confocal microscope.

#### CRISPR-mediated generation of FOXP2-mNeon line

gRNAs targeting within 50 base pairs (bp) of the FOXP2 STOP codon were designed using online tools CCTop (https://cctop.cos.uniheidelberg.de:8043/) and CRISPOR (http://crispor.tefor.net/). Four candidate gRNAs were selected based on high predicted specificity and efficiency scores, and a low number of off-target sites. Efficiency for each gRNA was determined by transfection with Cas9 into iPSCs as described below. A region across the FOXP2 stop codon was sanger sequenced and the resulting traces analysed using TIDE (http://shinyapps.datacurators.nl/tide/).The gRNA with the highest efficiency was chosen (AGAGCCTTTATCTGAAGATC).

The donor construct sequence contained sequences encoding a flexible XTEN linker^62^, a P2A peptide and mNeonGreen, surrounded by two homology arms, and was purchased as a double-stranded DNA gBlock (IDT). ssDNA copies of the repair construct were generated using the Guide-it™ Long ssDNA Production System v2 (Takara Bio, 632666). As prediction of the relative integration efficiency of each strand is unknown, both ssDNA products were made and used in separate transfection reactions.

A ribonucleoprotein was prepared by incubating 0.5 μl Alt-R S.p. HiFi Cas9 Nuclease V3 (IDT, 20925207), 22 pMole sgRNA and 1.5 μg of ssDNA in transfection enhancer buffer (Invitrogen, MPK10096) for 20 min at room temperature. For each reaction, a suspension of 2.2 x10^5^ cells was mixed with the prepared ribonucleoprotein and ssDNA donor solution. Cells were electroporated using the NEON Transfection system (Invitrogen, MPK10096) and 10 μl Neon pipette tips, according to program HiTrans (1400 V, 20 ms width, 1 pulse), and seeded onto Matrigel coated wells. gDNA was extracted from the resulting pools of transfected cells using the DNeasy Blood & Tissue Kit (Qiagen, 69504). To detect the desired insertion, PCR used primers outside the homology arm regions around the *FOXP2* STOP codon (Table S3) and products were run on a 1% agarose gel to visualise bands. Untransfected cells showed a clear band of ∼1.4 kb, whilst successful insertion generated a 2.2-kb band also present in the CRISPR pools.

To obtain stable reporter lines clones were isolated from the pool of transfected cells. The CRISPR pool was serially diluted onto mitotically inactivated CF1 mouse embryonic feeder cells (Millipore), allowing colony formation from single cells. Colonies were picked manually using a P200 tip, selected colonies were defined, relatively small and spaced apart from others to reduce the chance of cross contamination between clones. In total, 288 colonies were picked. iPSC clones were cultured until sufficiently confluent to be passaged to replica plates, allowing freezing of cell stocks and gDNA extraction. gDNA from 245 clones was screened for successful insertion of mNeon at the *FOXP2* locus by PCR using primers spanning from outside the 5’ homology arm to inside the mNeon sequence (Table S3). 75 clones produced a PCR product and used for the next screening step. Primers spanning outside the homology arms, as described for checking the pool, were used to genotype selected clones. 17 clones appeared homozygous for the insertion with only a higher molecular weight band around 2.2 kb, and 17 clones appeared heterozygous with both a 2.2-kb and WT 1.4-kb band. Sanger sequencing was performed on a selection of clones and verified correct sequences in two homozygous clones (A6B & B2D). These clones were expanded to create reporter line stocks to be used for following experiments and QC performed on this stock.

#### CRISPR clone quality control

Karyotype analysis: To test for gross chromosomal abnormalities, gDNA from each iPSC stock was analysed and compared to the parental stock. gDNA was extracted using the DNeasy Blood & Tissue Kit (Qiagen, 69504) and samples submitted for SNP karyotyping analysis (Infinium Global Screening Array-24 v3.0, Illumina, performed by Life&Brain GmbH).

Mycoplasma: iPSCs were tested for mycoplasma using the MycoAlert™ PLUS Mycoplasma Detection Kit (LT07-705).

Off-target site sequencing: The top six highest risk off-target sites were predicted for the gRNA by CCTop^63^ (https://cctop.cos.uniheidelberg.de:8043/) and CRISPOR^65^ (http://crispor.tefor.net/) and primers designed to amplify regions of approximately 200 bp around these high-risk sites (Table S3). Sanger sequencing was performed on gDNA from FOXP2-mNeon and compared to the parental AH017-3 stock.

Fluorescence-activated cell sorting (FACS) staining for pluripotency markers: iPSCs were harvested using TrypLE (Gibco, 12604013) and fixed in 2% PFA at room temperature for 10 mins. Cells were then centrifuged at 400 x g for 5 min, before resuspending in ice-cold methanol to permeabilise for at least 30 min at -20°C. Next, cells were washed once in FACS buffer (1% BSA in PBS) and incubated with primary antibodies against pluripotency markers TRA-1-60 and Nanog or isotope controls (Table S2) in FACS buffer at room temperature in the dark. After 1 hour of staining, cells were washed twice with FACS buffer and analysis performed using a Cytoflex LX analyser (Beckman Coulter).

Trilineage differentiation: To assess pluripotency of the iPSCs after CRISPR/Cas9 editing, a trilineage differentiation was performed according to the STEMdiff™ Trilineage Differentiation Kit (STEMCELL Technologies, 05230). After differentiation, coverslips were washed in PBS and fixed in 4% PFA for 20 min at room temperature. Fixed cells were permeabilised for 20 min using 0.4% Triton-X-100 in PBS and incubated in block (5% serum (normal goat serum, Abcam, ab7481 or normal donkey serum, Abcam, ab7475), 0.1% Triton-X-100 in PBS) for 1 hour. Primary antibodies (Table S2) in diluent (5% serum in PBS) were applied over night at 4°C. Following three washes in PBS, secondary antibodies were incubated for 1 hour at room temperature. Three washes were performed to remove unbound antibody with 1 μg/ml Hoechst (Invitrogen, H1398) in the final wash. Stained coverslips were imaged on a Zeiss-880 laser scanning confocal microscope.

#### Western blotting

To extract protein, cerebellar organoids were lysed with manual disruption in ice cold Pierce RIPA buffer (Thermo Scientific, 89900) supplemented with protease (cOmpleteTM, Roche, 04693159001) and phosphatase (PhosSTOPTM, Roche, 04906837001) inhibitors. Protein samples were then diluted with NuPAGE™ LDS Sample Buffer (Invitrogen, NP007) containing 25 mM DTT and denatured by heating at 70oC for 10 min. 30-50 μg of protein was loaded on a 4-12% gradient Bis-Tris gel (Invitrogen, NP0335) and run at 200 V for 50 min. Protein was then transferred to a nitrocellulose membrane over 1.5-2 hour at 250 mA. The membrane was blocked with 10% milk in TBST (150 mM NaCl, 10 mM Tris pH8, 0.05% Tween 20) and primary antibodies (Table S2) incubated overnight at 4°C, diluted in 3% BSA in TBST. After washing 3 times in TBST, secondary antibodies (HRP-conjugated, GE healthcare) were applied for 1-2 hour at room temperature in 3% BSA in TBST. Washing was repeated and then Pierce™ ECL Western Blotting Substrate (Thermo Scientific, 32209) was added to the membrane and signal captured using a Chemidoc MP imaging system.

#### Live-cell imaging

A live-cell z-stack of a whole FOXP2-mNeon organoid was taken using a Leica Mica confocal imaging system.

#### Isolation of mNeon-positive cell population

Organoids at day 63 of differentiation were washed in HHGN (1x HBSS (Invitrogen, 14180-046), 2.5 mM HEPES pH7.3-7.5 (Invitrogen 15630-049), 35 mM Glucose, 4 mM NaHCO3 (Sigma, S6297) in H2O) and incubated in neuronal isolation enzyme (Thermo Scientific, 88285) for 30-40 min at 37°C with resuspension every 10 min. After removal of enzyme, organoids were washed again in HHGN and gently dissociated to single cells in Differentiation medium by pipetting. Dissociated cells were centrifuged at 400 x g for 5 min and resuspended at 10^7^ cells/ml in FACS buffer (1% BSA (Sigma, A4161-5G) in PBS). To stain dead cells, 1 μg/ml DAPI (Biolegend, 422801) was added. Cells were sorted directly into RLT lysis buffer (Qiagen, 74004) using a BD FACSAria III (BD Biosciences).

#### RNA sequencing

RNA from sorted cell populations was isolated using the RNeasy Micro kit (Qiagen, 74004). RNA concentration was measured using the Quant-iT RiboGreen RNA kit (Invitrogen, R11490) and the RNA integrity number (RIN) evaluated using High Sensitivity RNA ScreenTape (Agilent, 5067-5579) on a TapeStation. All samples submitted for sequencing had a RIN of 8.9 or above.

Library preparation and sequencing was performed by the Oxford Genomics Centre. Library preparation used 22-58 ng of RNA per sample (NEBNext Ultra II Directional mRNA kit, New England BioLabs, E7760). Sequencing was performed an Illumina NovaSeq6000 platform using 150 bp directional paired end read sequencing. Read alignment and quantification was performed using Kallisto^64^ with Ensembl release 108. Over 45 million reads were aligned for all samples (45.7 – 52.5 million reads, 83-87% of total reads pseudoaligned).

Transcript quantifications were imported into R and mapped to genes by tximport (tximport_1.26.1) using a txtgene object (created with makeTxDbFromEnsembl from GenomicFeatures_1.50.3). Genes with more than 10 counts in 4 samples were included in analysis. DESeq2 (DESeq2_1.38.2, ^66^) was used for differential gene expression analysis using formula ∼Differentiation + mNeonpopulation. The lfcShrink function type ‘apeglm’ was applied to remove noise from low expressed genes ^67^. Genes with adjusted p-value (Benjamini-Hochberg method to correct for multiple testing) below 0.05 were determined significantly differentially expressed. For visualising, gene expression was normalised using variance stabilizing transformation (VST), additional plots were made using ggplot2 (ggplot2_3.4.0) and Pheatmap (pheatmap_1.0.12).

Gene set enrichment analysis (GSEA) was performed on VST normalised expression output from DESeq2 through the online tool (GSEA 20.4.0 on the GenePattern server https://www.genepattern.org/) ^68,69^. Analysis used 1000 repetitions, permutated on geneset and a false discovery threshold of 0.05 for significant terms. GO terms were used from the MSigDB on the GenePattern server, whilst cerebellar markers and disorder genesets were uploaded. Differentially expressed genes (absolute log fold change 1.5, adjusted p-value <0.05) from laser-capture micro-dissected fetal human cerebellar samples ^28^. Cerebellar cell-type marker genes extracted using scran::FindMarkers() from developing human snRNAseq ^26^. Disorder gene lists were previously used for analysis of human cerebellar data ^28^, with addition of ASD genes categorised as class 1 (High confidence) by Simons Foundation Autism Research Initiative (SFARI, https://gene.sfari.org/, 01-23-2023 release) and CAS genes compiled from publications that performed whole-genome sequencing of probands with CAS and close family members^54–56^. Genes differentially expressed in autopsy brain samples of individuals with different neurological disorders were published by Gandal et al., 2018^70^.

When comparing overlap with FOXP2 target genes^43–45^, the criteria for differentially expressed genes between mNeon-sorted organoid populations was an adjusted *p*-value <0.05 and an absolute log2FoldChange >1.5. GeneOverlap (GeneOverlap_1.34.0) was used to perform a Fisher’s exact test to determine significance of overlap.

## QUANTIFICATION AND STATISTICAL ANALYSIS

Statistical tests were performed in R. Significant differences between two groups were determined using student t-test. For more than two groups, ANOVA followed by post-hoc Tukey’s HSD test was used to identify significant differences. Unless overwise stated, a *p*-value threshold of 0.05 was used to determine significance and *p*-values are indicated as \**p*<0.05, \*\**p*<0.01, \*\*\**p*<0.001. Details of statistical tests are found in results and figure legends. Each biological replicate (n) resulted from one or more organoid(s) from an independent differentiation batch, therefore each experiment used three or more differentiations.

