## Supplemental Information for "Cerebellar organoids model cell type-specific *FOXP2* expression during human cerebellar development"

[1]Nuffield Department of Clinical Neurosciences, University of Oxford, Oxford OX3 9DU, UK
[2]Kavli Institute of Nanoscience Discovery, University of Oxford, Oxford OX1 3QU, UK
[3]Genome Engineering Oxford, Sir William Dunn School of Pathology, University of Oxford, Oxford OX1 3RE, UK
[4]James and Lillian Martin Centre for Stem Cell Research, Sir William Dunn School of Pathology, University of Oxford, Oxford OX1 3RE, UK
[5]Present address: Centre for Developmental Neurobiology, King's College London, London SE1 1UL, UK
[6]Lead contact

### Supplementary Methods

#### Analysis of mouse cerebellar samples

Wildtype mice (C57BL/6JOlaHsd) were sacrificed, and cerebella extracted under Home Office Project License PP4924146. Cerebellar samples were fixed in 4% PFA overnight at 4°C and then cryoprotected in 30% sucrose. Immunostaining was performed as for organoid samples.

#### Repair construct sequence for FOXP2-mNeon generation

```
GTATTTGTTTCAACAAACACCCAATCTAGCTTATACTGGAGAGTTTAAAAAAAAATAGAAATAAAATT
GTTAAACTCAAGTCTGAAATATTTAGAAATAACAGTTGTTATCTTAATTGCTAGCAAAAAACTGAATG
GTAAAAGTGTTTTCATTTGTTGATTAATTATTTGTTTACTATTCCTTTACTACCACTCCTTGATTATC
ATGTTATAACAAATACTAACAAAACATTTCATTCAGAAAGAATGCTGAGAAACGGATGTTTTACCTAA
ACAGCACAGCATGTGGGAGTCCATGCTCACATTACTCTGAATCCCTGCCTCCACAGTTCTACTGTAAA
GTATCATGTCTCTAACCAGCTCACGCAATCAATCAGATGCCCTTTTAAGAAAGACAATGTTGTGTCTT
CTTGCCTTTTTTCAGTTGACCTCTTCACTGCAAAGTTGGCCAAACTCTGTTTGTTGCTTACTTAGTAA
AATTTTGGTGTATCTACATGTTTTTCAGACATTCAATCCACGTCAAGGAAGAGCCAGTGATTGCAGAG
GATGAAGACTGCCCAATGTCCTTAGTGACAACAGCTAATCACAGTCCAGAATTAGAAGACGACAGAGA
GATTGAAGAAGAGCCTTTATCTGAAGATCTCGAATCCGGTAGCGAAACACCGGGGACTTCAGAATCGG
CCACCCCGGAGTCTGCCACGAACTTCTCTCTGTTAAAGCAAGCAGGAGACGTGGAAGAAAACCCCGGT
CCTATGGTGAGCAAGGGCGAGGAGGATAACATGGCCTCTCTCCCAGCGACACATGAGTTACACATCTT
TGGCTCCATCAACGGTGTGGACTTTGACATGGTGGGTCAGGGCACCGGCAATCCAAATGATGGTTATG
AGGAGTTAAACCTGAAGTCCACCAAGGGTGACCTCCAGTTCTCCCCCTGGATTCTGGTCCCTCATATC
GGGTATGGCTTCCATCAGTACCTGCCCTACCCTGACGGGATGTCGCCTTTCCAGGCCGCCATGGTAGA
TGGCTCCGGATACCAAGTCCATCGCACAATGCAGTTTGAAGATGGTGCCTCCCTTACTGTTAACTACC
GCTACACCTACGAGGGAAGCCACATCAAAGGAGAGGCCCAGGTGAAGGGGACTGGTTTCCCTGCTGAC
GGTCCTGTGATGACCAACTCGCTGACCGCTGCGGACTGGTGCAGGTCGAAGAAGACTTACCCCAACGA
CAAAACCATCATCAGTACCTTTAAGTGGAGTTACACCACTGGAAATGGCAAGCGCTACCGGAGCACTG
CGCGGACCACCTACACCTTTGCCAAGCCAATGGCGGCTAACTATCTGAAGAACCAGCCGATGTACGTG
TTCCGTAAGACGGAGCTCAAGCACTCCAAGACCGAGCTCAACTTCAAGGAGTGGCAAAAGGCCTTTAC
CGATGTGATGGGCATGGACGAGCTGTACAAGTGAGAACTGACTTGTGAAACGTCAGCGTGAAGGGACA
TATCACTGACCTTCATAACCACTCCACAACCATGAATATTTGACAAATTTTTACTGTGACTATTTATT
AAGCATGGATAAAGGAGACAGCCCTAAAGGAACTTACTAAGCCAGCCCTTTGGGATTCAGTACCAACA
GGCAAATTGCTTGTTTTCTTCTTCTTCTTCTTTTTTTTTTTAGAAAAAAAGACAAAAACTGATTT
TCTTGAAAAAAAAAAATGAACTGTTCTTTCTATAATGGCTTTGCCCATTTAAAAAATGTGGCTCTTAA
GGGTTCATGAAATGACTGAATATGAGGATACATGTCCTGTAGAAAGCAAATGCGCCTCATATACTGCC
AAAAATAGTGTTAGTTTCATTAATGTGAATTTTCCAGCATTCAGTAGTTGTAATGTTAGAAACAATTG
CTGGTCAAGTTCAACTTGTTGCTATTGTTTTTAATTTGCACAGGAGTAGTATCAGAAATTAGTGTCAC
TGCTTGTATCTAGCTGAATTTTAAACAACAGAACATTAGTTTTTTATGTTGGTGCCACCAACTGTAAA
TGACATAAGTTAGTTATTACAAAACACAGTAATTAGACTGTTGCAACCATCTAAAACCTTAGGCTTCC
AGTCTGTGCTG
```

**Table S1. RT-qPCR primers**

| Target | Fw primer | Rv primer |
|---|---|---|
| *ACTIN* | GCCGCCAGCTCACCATGGATG | CCATCACGCCCTGGTGCCTGG |
| *ATOH1* | TGTTATCCCGTCGTTCAACAAG | TGGGCGTTTGTAGCAGCTC |
| *EN1* | GCTTGTCCTCCTTCTCGTTC | TGGTCAAAACTGACTCGCAG |
| *EN2* | CCGGCGTGGGTCTACTGTA | GGCCGCTTGTCCTCTTTGTT |
| *FOXP1* | CTTCAGCTTCCTCTGGATCG | GGCAGATCTCCTATGCAAGC |
| *FOXP2* | TGGCAGCAGAGATGGAAGAT | AGTTGTCTTGCTGCCTGGAG |
| *FOXP4* | AATGGTGAGATGAGTCCCGC | TTTGCTGTCATTGTTCCCTGG |
| *GAPDH* | GGAAGGTGAAGGTCGGAGTC | GTTGAGGTCAATGAAGGGGTC |
| *GBX2* | AAAGAGGGCTCGCTGCTC | GGTCGTCTTCCACCTTTGAC |
| *SKOR2* | AGCCCAGTTCACCATCCAT | GCTGTTGTCATCCTTTGTAGATAC |

**Table S2. Primary antibodies**

| Target | Supplier | Code | Source | Use | Concentration |
|---|---|---|---|---|---|
| Actin | Abcam | Ab179467 | Rabbit | WB | 1:1000 |
| Brachyury | R&D systems | AF2085 | Goat | IF | 1:100 |
| CALB1 | Swant | CB38 | Rabbit | IF | 1:10,000 |
| FOXA2 | R&D systems | AF2400 | Goat | IF | 1:200 |
| FOXP1 | CST | 2005 | Rabbit | IF | 1:500 |
| FOXP2 | Abcam | Ab1307 | Goat | IF | 1:1000 |
| FOXP2 | Sigma | MABE415 | Mouse | WB | 1:1000 |
| FOXP4 | Atlas | HPA007176 | Rabbit | IF | 1:1000 |
| GBX2 | ThermoFisher | H00002637-M01 | Mouse | IF | 1:100 |
| IgG 647 | Cell Signalling | 2985S | Rabbit | FACS | 5 µg/ml |
| IgM 488 | Biolegend | 401617 | Mouse | FACS | 15 µg/ml |
| KIRREL2 | R&D | Af2930 | Goat | IF | 1:250 |
| MAP2 | Abcam | Ab183830 | Rabbit | IF | 1:500 |
| mNeonGreen | Chromotek | 32F6 | Mouse | IF | 1:1000 |
| mNeonGreen | Chromotek | 4M2 | Rabbit | WB | 1:1000 |
| NANOG 647 | Cell Signalling | 5448S | Rabbit | FACS | 5 µg/ml |
| NCAM | Abcam | ab75813 | Rabbit | IF | 1:200 |
| NESTIN | Sigma | MAB5326 | Mouse | IF | 1:500 |
| PAX6 | GeneTex | GTX113241 | Rabbit | IF | 1:500 |
| SKOR2 | Atlas antibodies | HPA046206 | Rabbit | IF | 1:500 |
| SOX17 | R&D systems | AF1924 | Goat | IF | 1:100 |
| TRA-1-60 488 | Biolegend | 330614 | Mouse | FACS | 15 µg/ml |
| ZFHX4 | Atlas antibodies | HPA023837 | Rabbit | IF | 1:500 |

### Table S3. PCR primers used in FOXP2-mNeon screening and quality control

| Purpose | Name | Fw primer | Rv primer | Product size (bp) |
|---|---|---|---|---|
| Sequencing *FOXP2* homology arms Clone screening: genotyping | FOXP2_HAseq | TTGAAAATAGTACTGGCATCCTGG | CTATGGTGCTTGCACAGATAATG | WT *FOXP2*: 1438 FOXP2-P2A-mNeon: 2245 |
| Additional *FOXP2* sequencing | FOXP2_HAs_midse2 | GCCAGTGATTGCAGAGGATG | - | NA |
| Additional *FOXP2* sequencing | FOXP2_HAseq_midseq_rv | - | TTTGGCAGTATATGAGGCGC | NA |
| FOXP2 gRNA testing | FOXP2_gRNA_test | CAATCCACGTCAAGGAAGAGC | TGCCTGTTGGTACTGAATCC | 314 |
| Generation of P2A-mNeon ssDNA repair construct | FOXP2_ssDNA | TTCAACAAACACCCAATCTAGC | CAGCACAGACTGGAAGCCTA | 2221 |
| Clone screening insertion | FOXP2mNEON_screen | TGGGGTTGGAGGGATAATAAACTT | CCATCATTTGGATTGCCGGT | WT *FOXP2*: no product *FOXP2-P2A-mNeon*: 1081 |
| Off-target testing | 10_OTUD1 | AAGAGCTCTGAAGCCTGCTG | AAGTGTGGGGATTACAGGCG | 196 |
| Off-target testing | 8_KBTBD11 | GCACCATTGTACTCCAGCCT | AGCGGAACACAATAAATGTGGA | 133 |
| Off-target testing | 10_PDZD8 | GCGTTACTGTTGTGGGAGGA | GCCCTTAGAGCTGTCTGCAT | 231 |
| Off-target testing | 6_COX7A2 | GTCCGCTGAGAGTTTCCTCC | TTTGGTTCAGAAAGGCGGGA | 506 |
| Off-target testing | X_RP1 | GAGTACACCCAAGGCCTGATT | ACACATATACCTTCCCAACACACT | 188 |
| Off-target testing | 3_ACPP | ACGTTGACCGGACTTTGATG | ACAACTGTTGAGTAGCCTGC | 206 |

### Supplementary Figures

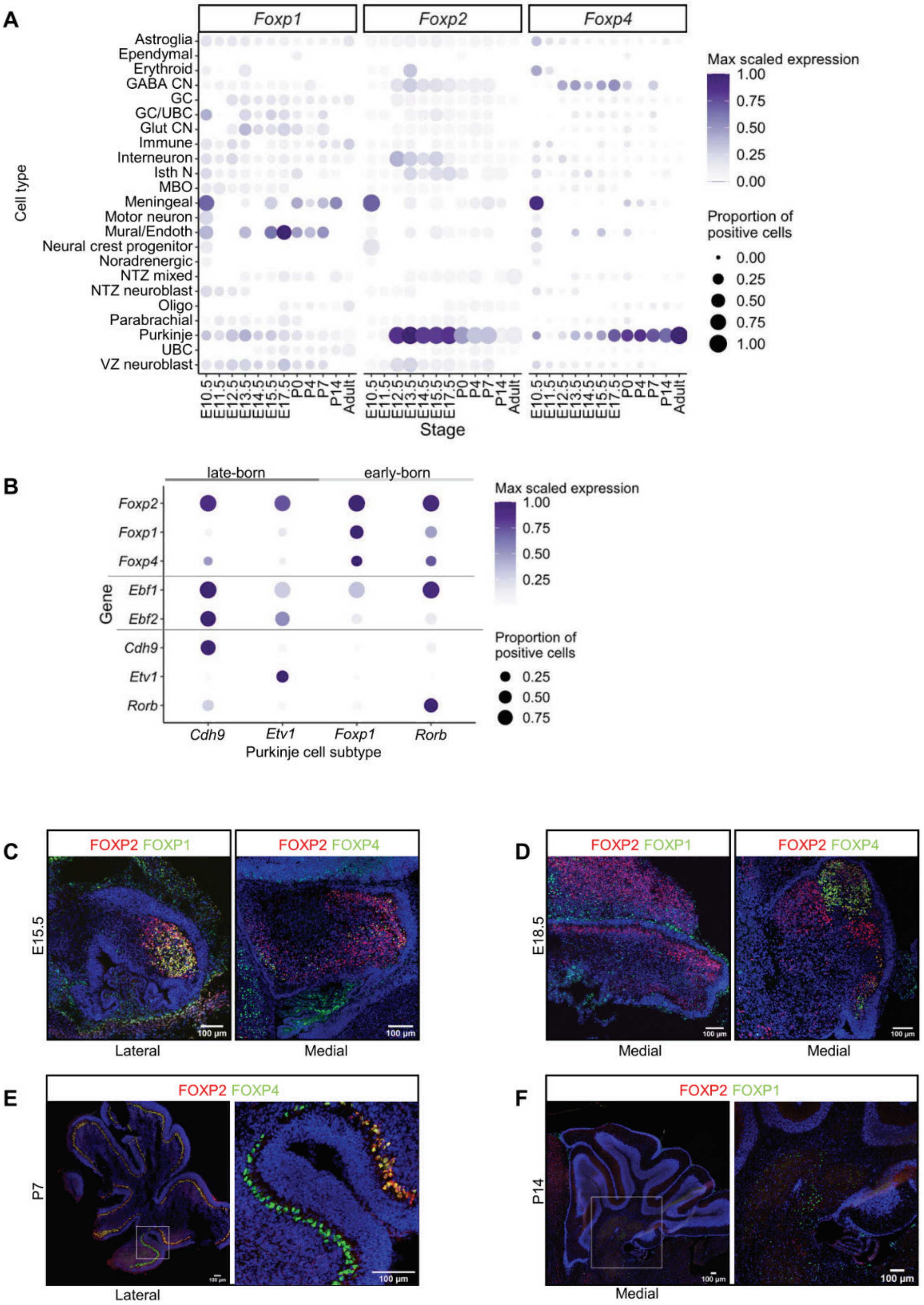


**Figure S1. *Foxp* family genes are expressed in developing mouse Purkinje cells.**
**(A)** Expression of *Foxp* genes in the developing mouse cerebellum, plotted across timepoints and cell types based on published snRNA-seq data from Sepp et al., 2024. Point size indicates the proportion of expressing cells in each cluster and the colour scale shows average expression levels. Samples with <10 cells were excluded. (B) *Foxp* genes show different expression patterns across mouse embryonic Purkinje cell subtypes. Purkinje cell subtypes can be distinguished by levels of *Ebf1* and *Ebf2*, and high expression of markers *Cdh9*, *Etv1*,

*Foxp1* and *Rorb*. Analysis based on snRNAseq data from Sepp et al., 2024. **(C-F)** Immmunostaining for FOXP proteins FOXP2 (red) with FOXP1 or FOXP4 (green) across mouse cerebellar development confirm expression in developing Purkinje cells. FOXP1 and FOXP4 display differential expression across Purkinje cell clusters. In addition, FOXP1 and FOXP2 show sparse expression in the white matter of post-natal samples. Nuclei labelled by Hoechst (blue), scale bars = 100 µm. Insert locations indicated by white box in E&F.

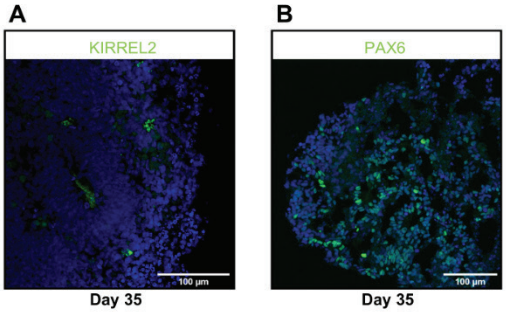


**Figure S2. Cerebellar organoids generate cells from both germinal zones.**
Immunostaining of day 35 cerebellar organoids using antibodies against **(A)** KIRREL2 labelling the ventricular zone and **(B)** PAX6 labelling the rhombic lip lineage. Nuclei labelled by Hoechst (blue), scale bars = 100 µm.

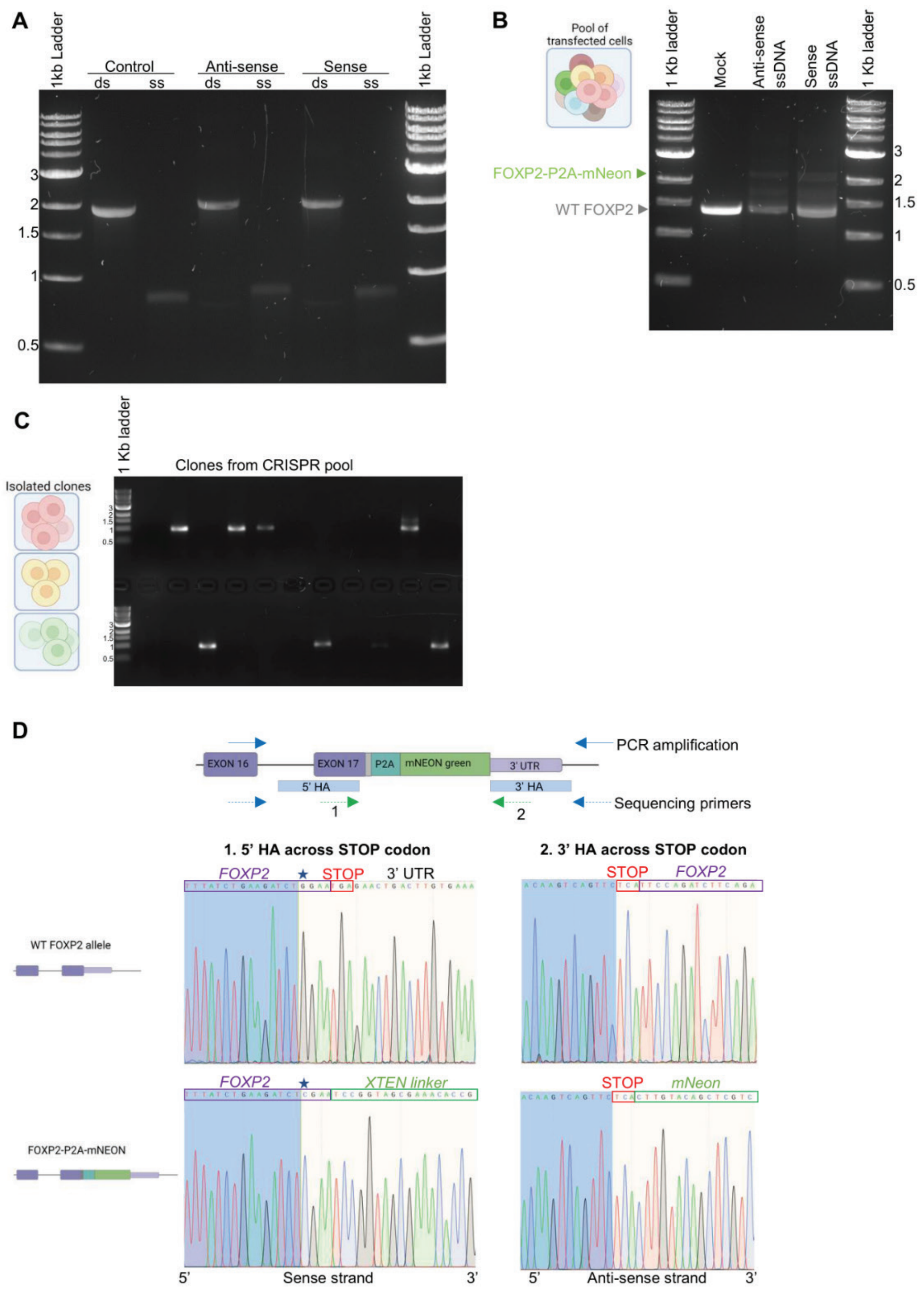


**Figure S3. CRISPR/Cas9-mediated generation of FOXP2-mNeon reporter line.**
**(A)** The size and purity of the ssDNA donor construct was checked before transfection. ssDNA products run at approximately half the size of the parental dsDNA (2.1 kilobase pairs (kb)) and show clean bands indicating full digestion. **(B)** Pools of transfected cells show insertion of P2A-mNeon when tested by PCR and visualised on a 1% agarose gel. From untransfected cells, the PCR generated a wild-type (WT) band ~1.4 kb. Transfections with either ssDNA donor resulted in some cells with successful insertion detected by an additional band at ~2.2 kb, with similar efficiency. **(C)** To screen clones for insertion, a primer pair from outside the 5' homology region to within the mNeon sequence were used. PCR products were visualised on a 1% agarose gel, with the presence of a band showing mNeon insertion. **(D)** Sanger sequencing of the genotyping PCR product in C confirmed the accurate insertion of the mNeon.

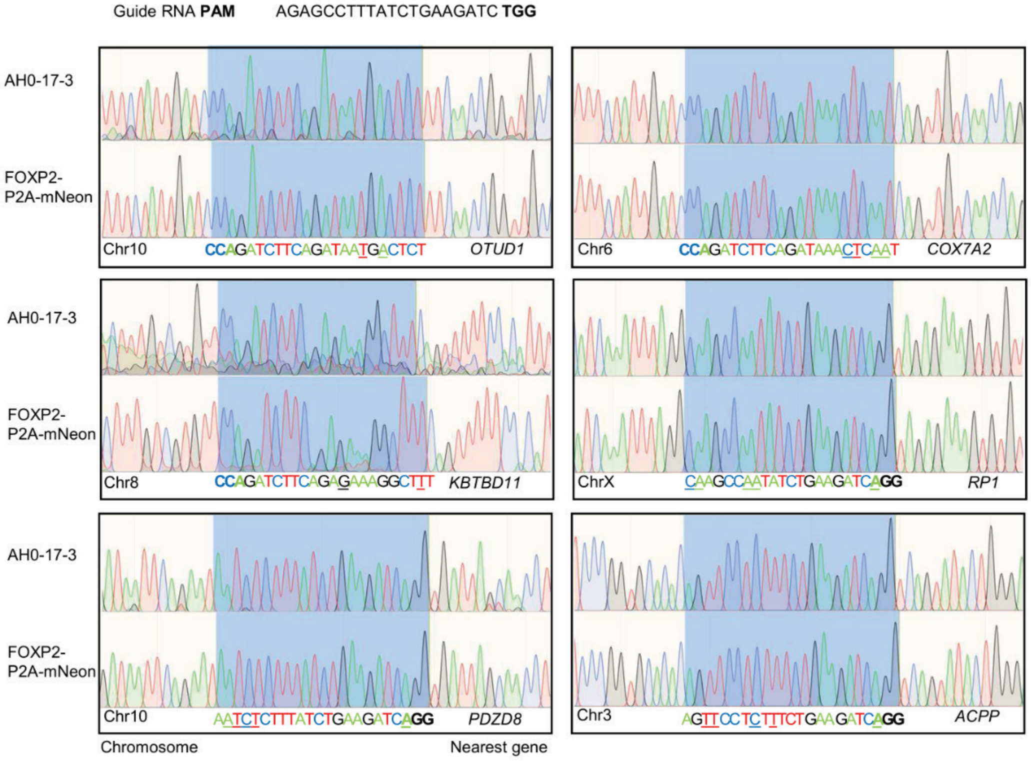


**Figure S4. The sequences of predicted off-target sites were not altered in the FOXP2-mNeon line.**

The predicted top six off-target sites for the gRNA used for FOXP2 editing were sequenced to check for mutations in the FOXP2-mNeon line. Sequencing traces are shown for the parental AH017-3 line and FOXP2-mNeon reporter line clone B2D. Sites with a close match to the gRNA target sequence and their adjacent PAM motif are highlighted in blue. Sequences are shown below with mismatches to gRNA target site underlined and the PAM sequence in bold. The off-target loci are labelled with their chromosome and nearest gene.

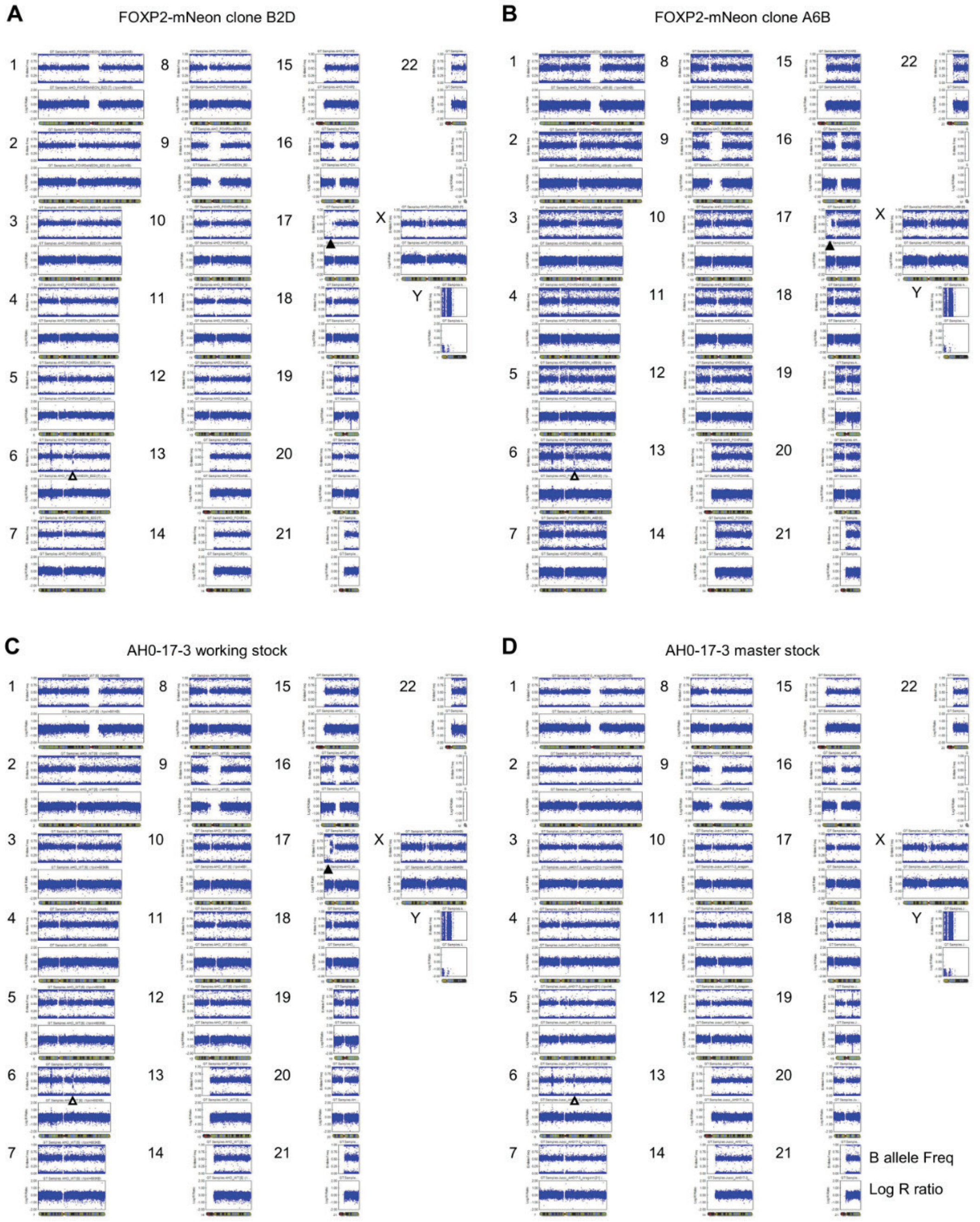


**Figure S5. Karyograms from SNP analysis of FOXP2-mNeon clones and parental iPSC line.**
SNP array analysis on genomic DNA was used to assemble karyograms for the iPSC stocks. Karyotypes are shown for two FOXP2-mNEON reporter line clones **(A)** B2D & **(B)** A6B, **(C)** the parental AH017-3 working stock used to generate this line and **(D)** an alternative AH017-3 master stock. The control iPSC line, AH017-3, had a small duplication when established (Chr 6q14.1 2mb duplication) indicated by the open triangle in all 4 samples (Handel *et al.,* 2016). However, the AH017-3 stock used for reporter line generation shows an additional chromosomal alteration on chromosome 17 which was consequently shared in both FOXP2-P2A-mNeon clones. The solid triangle indicates the Chr17p13-12 13mb loss of heterozygosity (LOH) and Chr17p12-11 6mb duplication.

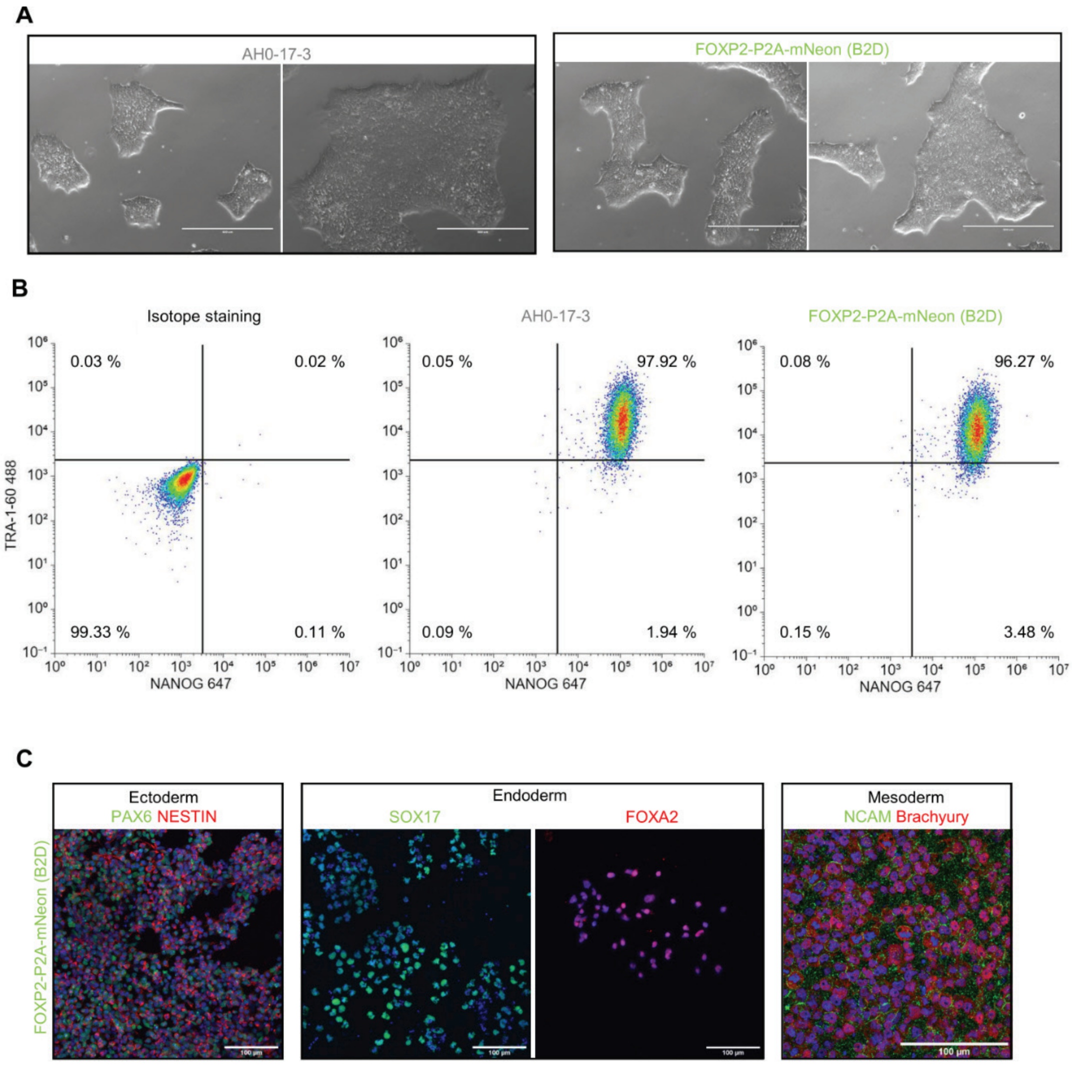


**Figure S6. FOXP2-mNeon line clone B2D displays pluripotent state.**

**(A)** Brightfield images of control parental line (AH017-3) and FOXP2-mNeon reporter line (clone B2D) showing iPSC colonies with bright borders and tightly packed cells with large nuclei. Scale bars = 400 µm. **(B)** Flow cytometry analysis was used to quantify expression of pluripotency markers. iPSCs were stained with fluorescent-conjugated antibodies for TRA-1-$60^{488}$ and $NANOG^{647}$ or with isotype controls $IgM^{488}$ and $IgG^{647}$ to set gating for positive test antibody staining. The population percentages for each quadrant are indicated. **(C)** To confirm the capability to differentiate into all three germ lineages, a trilineage differentiation was performed from FOXP2-mNeon iPSCs (clone B2D). Differentiated cells were fixed and stained using antibodies against markers for each lineage. Ectoderm cells were stained with antibodies against PAX6 (green) and NESTIN (red), endoderm cells separately with SOX17 (green) and FOXA2 (red), and mesoderm cells with NCAM (green) and Brachyury (red). Nuclei stained with Hoechst (blue), scale bars = 100 µm.

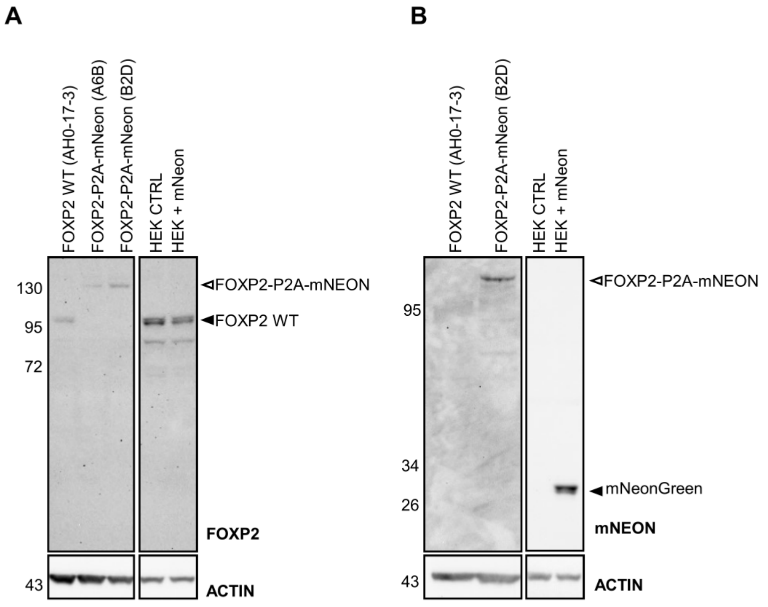


**Figure S7. FOXP2-mNeon line shows incomplete cleavage of FOXP2 and mNEON.**
**(A)** Western blotting for FOXP2 shows a higher molecular weight band in two FOXP2-mNeon reporter line clones (A6B & B2D) at ~130 kDa, the predicted size of a FOXP2-mNeon fusion protein. A wild-type (WT) band for FOXP2 at around 95 kDa is present in the WT (AH017-3) cerebellar organoid and HEK293 cell samples. Protein samples were lysed from day 63 cerebellar organoids. ACTIN is shown as a loading control. **(B)** Western blotting for mNeonGreen shows a high molecular weight band in two FOXP2-mNeon reporter line clones (A6B & B2D) at ~ 130 kDa, the predicted size of a FOXP2-mNeon fusion protein. A band at 26kDa is detected in HEK293 cells with exogenous expression of mNeon. Samples are shown from HEK293 cells that were untransfected (HEK CTRL) or transfected with a plasmid to express mNeongreen (HEK+mNEON). ACTIN is shown as a loading control.

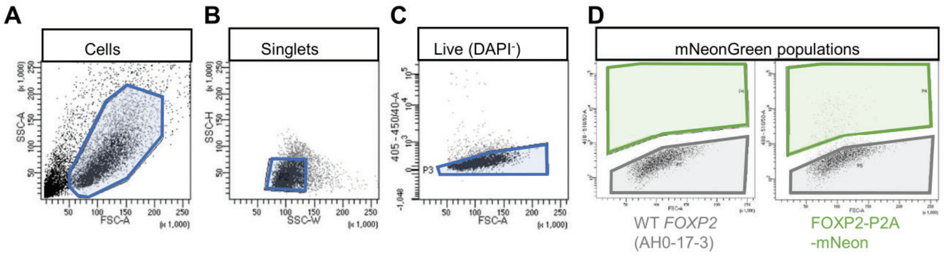


**Figure S8. Isolation of mNeon positive and negative cells, by flow cytometry.**
**(A)** Cells were selected from debris and other particles using a plot of forward scatter (x axis) against side scatter (y axis). **(B)** Single cells were then selected based on side scatter height (x axis) and side scatter width (y axis). **(C)** Live cells were selected based on low signal for DNA binding dye DAPI indicating intact membranes. Fluorescence intensity from 405 laser excitation on the y axis. **(D)** Finally, mNeon positive and negative populations were selected according to fluorescent intensity (y axis). Gating thresholds were determined using WT cerebellar organoids (left) which had no mNeon expression.

A

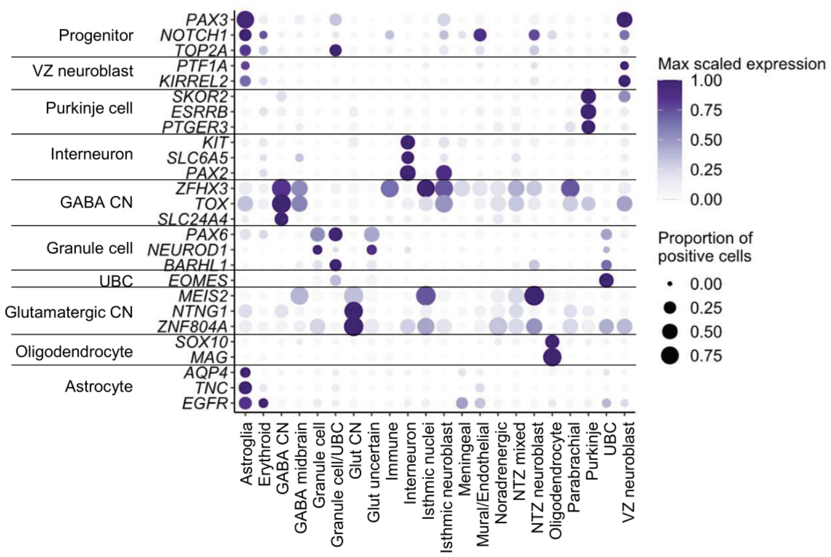


**Figure S9. Expression of selected cerebellar cell type markers show specificity in human snRNAseq.**
Expression of selected cerebellar cell types markers in published developing human snRNAseq (Sepp et al., 2024). Point size indicates the proportion of expressing cells in each cluster and the colour scale shows average expression levels. Samples with <10 cells were excluded. CN, cerebellar nuclei neurons; glut: glutamatergic; NTZ; nuclear transitory zone; UBC: unipolar brush cells; VZ: ventricular zone.
